# Monogamy promotes worker sterility in insect societies

**DOI:** 10.1101/059154

**Authors:** Nicholas G. Davies, Andy Gardner

**Affiliations:** Department of Zoology, University of Oxford, United Kingdom.; School of Biology, University of St Andrews, United Kingdom

**Keywords:** eusociality, haplodiploidy, Hymenoptera, inclusive fitness, kin selection, population genetics, promiscuity, worker non-reproduction

## Abstract

Inclusive-fitness theory highlights monogamy as a key driver of altruistic sib-rearing. Accordingly, monogamy should promote the evolution of worker sterility in social insects when sterile workers make for better helpers. However, a recent population-genetics analysis (Olejarz *et al.* 2015) found no clear effect of monogamy on worker sterility. Here, we revisit this analysis. First, we relax genetic assumptions, considering not only alleles of extreme effect—encoding either no sterility or complete sterility—but also alleles with intermediate worker-sterility effects. Second, we broaden the stability analysis—which focused on the invasibility of populations where either all workers are fully-sterile or all workers are fully-reproductive—to identify where intermediate pure or mixed evolutionarily-stable states may occur. Finally, we consider additional, demographically-explicit ecological scenarios relevant to worker non-reproduction. This extended analysis demonstrates that an exact population-genetics approach strongly supports the prediction of inclusive-fitness theory that monogamy promotes sib-directed altruism in social insects.

## Introduction

Altruism among animals is epitomised by the workers of insect societies, who sacrifice their personal reproductive success to promote their siblings′ welfare. This remarkable self-abnegation— seemingly at odds with the “survival of the fittest”—is traditionally explained by kin selection: a gene causing workers to share provisions or defend the communal nest can spread if the workers′ sacrifice increases the survival of their siblings, who are likely to carry copies of the same gene. Higher genetic relatedness between the altruist and her beneficiaries would therefore—all else being equal—promote selection for altruism (Hamilton 1964). Accordingly, monogamy is often highlighted as a key promoter of sibling altruism, since maternal promiscuity decreases relatedness between siblings, diminishing the inclusive-fitness benefits of sib-rearing (Hamilton 1972; Charlesworth 1978; Charnov 1978; Boomsma 2007, 2009, 2013; Gardner et al. 2012; Davies et al. 2016). A wealth of empirical evidence supports this view, revealing a strong association between monogamy and sib-directed altruism in arthropods (Hughes *et al.* 2008), birds (Cornwallis *et al.* 2010), and mammals (Lukas & Clutton-Brock 2012).

A conspicuous example of sib-directed altruism in the social Hymenoptera (wasps, bees, and ants) is worker sterility. in many hymenopteran species, female workers lay unfertilised eggs in their natal colony, which develop into males on account of their haplodiploid mode of sex determination. But in some species, workers have partly or entirely stopped making sons to focus their efforts on helping instead. A standard account of inclusive-fitness theory would predict that—as with other forms of sibling altruism—monogamy should promote helpful worker sterility.

However, this prediction has recently been challenged by Olejarz et al.’s (2015) mathematical analysis of worker sterility in haplodiploid insect colonies, which uses an intricate population-genetics model to derive exact conditions for the invasion and stability of a worker-sterility allele. Surprisingly, this analysis could not identify a consistent effect of monogamy on the evolution of non-reproductive workers. In this *Research Advance*, we revisit this analysis, exploring alternative assumptions concerning the genetics, evolution, and ecology of worker sterility. We find that a more-comprehensive investigation of Olejarz *et al.*’s (2015) exact population-genetics approach strongly supports the view that monogamy promotes helpful worker sterility in insect societies and corroborates inclusive-fitness theory more generally.

## Unconstrained allelic effects: monogamy promotes worker sterility

Olejarz *et al.* (2015) investigated the spread of an allele that renders workers carrying the allele— who would otherwise produce sons through arrhenotokous parthenogenesis, substituting them for the queen′s sons—completely sterile. As the proportion *z* of sterile workers in a colony increases, the proportion *p_z_* of males produced by the queen rather than by workers also increases, while overall colony productivity *r_z_* may increase or decrease. Following these assumptions, they found that—in a seeming challenge to inclusive-fitness theory—worker sterility sometimes invades under single mating (*n* = 1) only, sometimes under double mating (*n* = 2) only, sometimes under both single and double mating, and sometimes under neither, suggesting no clear effect of monogamy on the invasion of sterility (Olejarz *et al.* 2015).

To explore the generality of this unexpected finding, we take up a suggestion by Olejarz *et al.* (2015, p. 13) and extend their analysis to consider alleles with intermediate effects on worker sterility (as was done for a similar model by Olejarz *et al.* 2016). Intermediate-effect alleles may exhibit incomplete penetrance (such that each carrier has an intermediate probability of being sterile), or may encode intermediate phenotypes (such that each carrier divides her resources between colony tasks and personal reproduction); these scenarios are mathematically equivalent, but for simplicity, we focus on the former. This suggested extension seems particularly apt, as the incomplete penetrance of sterility has been shown to be important for the evolution of reduced worker reproduction both in theory and in empirical practice (Charlesworth 1978; Ratnieks *et al.* 2006; Wenseleers & Ratnieks 2006b; Ronai *et al.* 2016); indeed, some form of incomplete penetrance is required to preserve the fecundity of queens carrying the sterility allele. Accordingly, we have derived exact conditions for the invasion of a recessive or dominant sterility allele with arbitrary penetrance (see Methods). When we require mutant worker-sterility alleles to show full penetrance, our analysis exactly recovers Olejarz *et al*.’s (2015) results (Fig. 1a). However, when we allow mutant worker-sterility alleles to show incomplete penetrance, we find that—strikingly—monogamy always promotes the invasion of helpful worker sterility (Fig. 1b). (Note that monogamy may inhibit worker sterility when sterility is harmful; see Methods.)

Why does allowing intermediate effects make such a categorical difference? The population genetics of invasion is the key. For example, a recessive sterility allele, when rare, is almost always expressed in colonies founded by a heterozygous female who has mated with one mutant male and *n* − 1 wild-type males. Other colony types occur, but are either comparatively rare (because they require more copies of the mutant allele among mating partners), or exhibit exactly the same phenotype as wild-type colonies (because sterility is expressed only when both parents pass the recessive mutant allele to their daughters). Therefore, sterility can only invade if these “mutant” colonies—in which a proportion 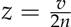 of workers are sterile, where *v* is the allele′s penetrance— succeed in spreading the sterility allele. If we only permit alleles with full penetrance (*v* = 1) arise, this allelic constraint may overpower the altruism-promoting effect of higher relatedness: for example, double mating (*n* = 2) may facilitate sterility′s invasion over single mating (*n* = 1) if colony efficiency is relatively high when *z* = 1/4 and relatively low when *z* = 1/2 (Fig. 1c). In contrast, if we permit alleles with incomplete penetrance (0 < *v* ≤ 1) to arise, mutant colonies may exhibit any one of a range of phenotypes, depending on *v* (namely, 0 < *z* ≤ ≤ 1/2 for single mating, and 0 < *z* < ≤ 1/4 for double mating), and monogamy always promotes the invasion of helpful worker sterility over promiscuity, by both maximizing sibling relatedness and allowing a wider range of phenotypes to be explored (Fig. 1d; see Methods for the corresponding analysis assuming dominant sterility).

**Figure 1:**
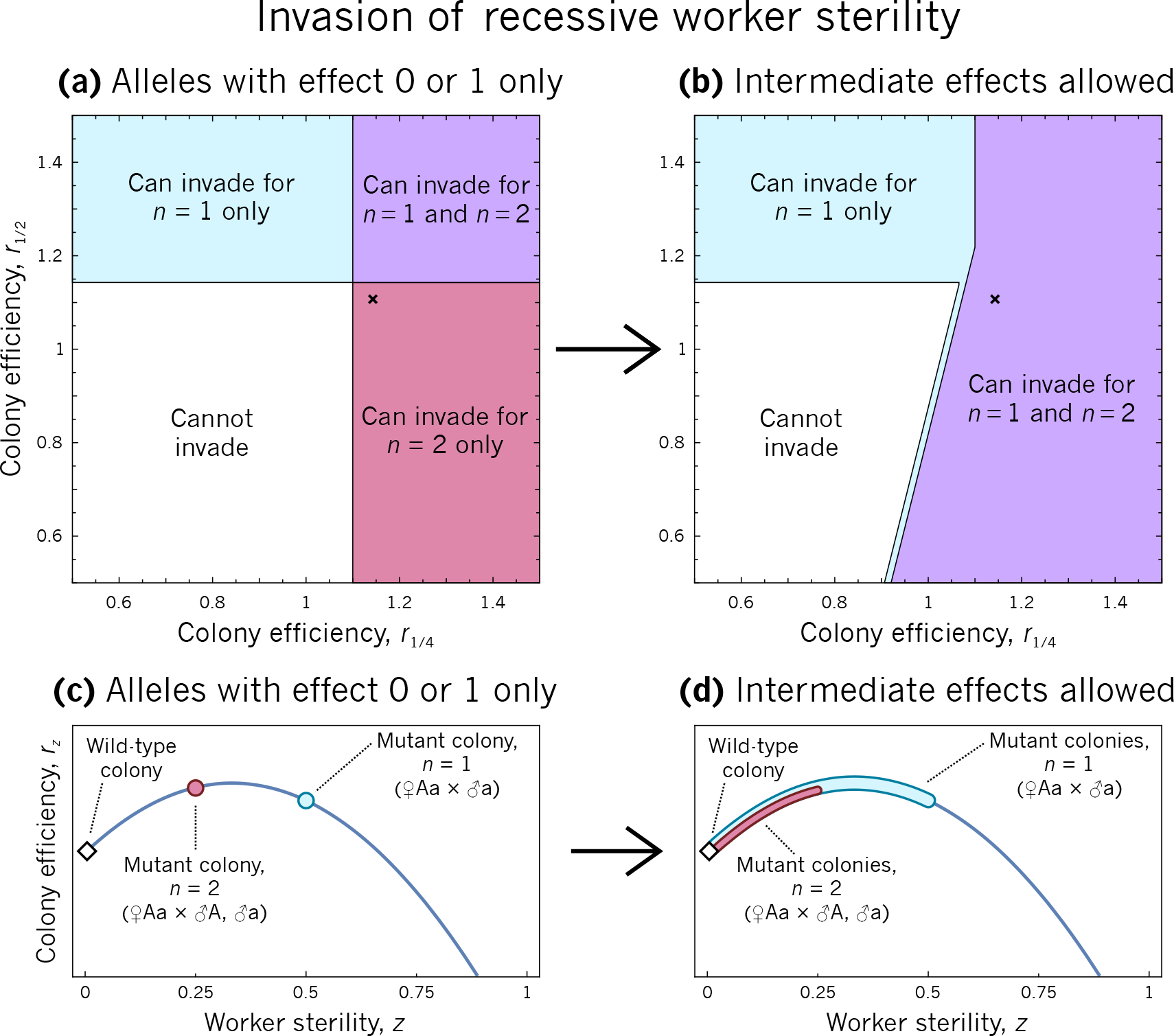
The invasion of worker sterility under recessive genetics, exploring the regions of parameter space where sterility can invade under single mating only, double mating only, both, or neither. **(a)** If we assume that only full-sterility alleles can arise, double mating sometimes promotes the invasion of sterility over single mating. But **(b)** if we assume that alleles encoding intermediate worker sterility may arise, double mating never promotes the invasion of sterility over single mating, depending on the colony efficiency values *r*_0_ = 1, *r*_1/4_, and *r*_1/2_. This is because **(c)** for a rare allele encoding full sterility, mutant colonies have the phenotype z = 1/2 under single mating and z = 1/4 under double mating. Therefore, sterility may invade more easily under double mating if colony efficiency is relatively peaked near z = 1/4. But **(d)** for a rare allele encoding intermediate sterility, mutant colonies may express any phenotype 0 < z ≤ 1/2 under single mating and 0 < z ≤ 1/4 under double mating, depending on the allele′s effect, and so mutant phenotypes are less constrained by the population′s mating number. In order to facilitate comparison with Fig. 3A of Olejarz *et al.* (2015), we assume *p_z_* = 0.2 + 0.8z, and for *r_z_* we use the unique quadratic curve passing through the points specified by *r*_0_ = 1, *r*_1/4_ and *r*_1/2_.

## Beyond invasion: monogamy promotes worker sterility

These results explain why promiscuity sometimes promotes the invasion of helpful sterility over monogamy under specific genetic constraints. But to only consider whether sterility invades may be misleading, for two reasons. First, that a sterility allele spreads from rarity says little about its equilibrium frequency, which may be a more-relevant measure of monogamy′s impact on worker altruism than mere invasion. Indeed, although promiscuity sometimes promotes sterility′s invasion *per se* under constrained penetrance, we find that monogamy typically promotes equilibrium sterility under the same conditions (Fig. 2).

Second, if we do allow intermediate-effect alleles, then considering only a single invasion is inadequate, because long-term evolution is likely to involve multiple successive invasions (*cf.* Ham-merstein 1996). How can we predict the outcome without knowing in advance which alleles may arise, and when? The solution is that, over the long term, populations exposed to sufficient genetic variation will converge on an evolutionarily-stable strategy (ESS; Maynard Smith & Price 1973)—a level of sterility that cannot be invaded by an allele encoding any other level of sterility. To identify a candidate ESS for sterility, we further extend *Olejarz et al.′s* (2015) population-genetics analysis to derive an exact condition for the invasion of an allele encoding a small increase to average sterility, z:

**Figure 2:**
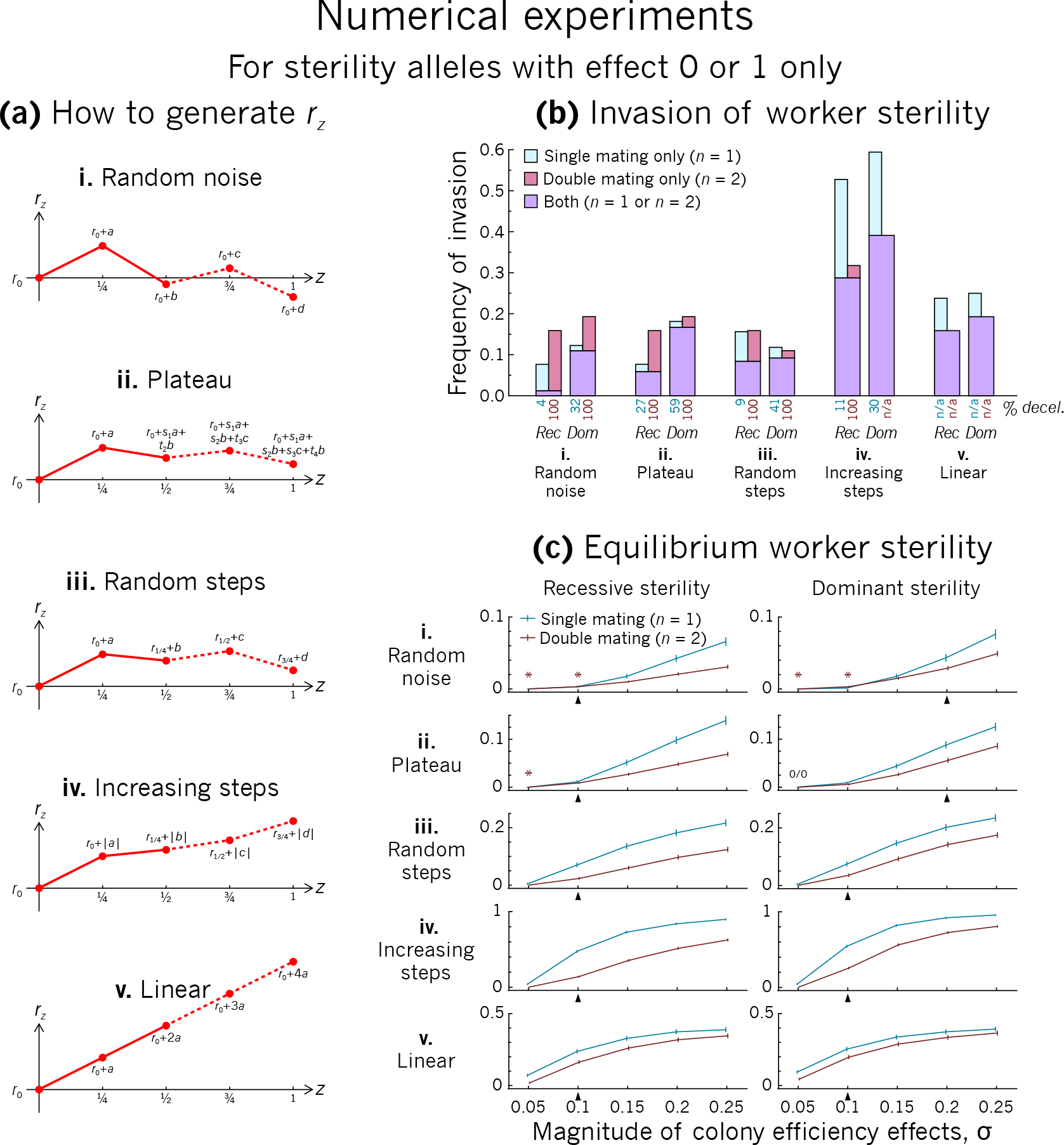
Here, we compare the evolution of worker sterility under single versus double mating by revisiting the numerical experiments of Olejarz et al. (2015). **(a)** There are many possible ways to construct the colony efficiency function r_z_ based on picking random numbers from a normal distribution. Five alternatives are shown here, including the two procedures used by Olejarz et al. (“Random noise”, their Procedure 1, and “Plateau”, their Procedure 2). For testing whether sterility invades, only two points are needed (solid lines), but this can be extended to four points (dashed lines) for measuring sterility at equilibrium. **(b)** We record the frequency of invasion of a full-sterility allele under single (*n* = 1) versus double mating (*n* = 2), running 10 million experiments for each scenario. Percentages beneath the bar chart show that an initially-decelerating r_z_ is required for sterility to invade under double mating only (see Methods). **(c)** We record the average worker sterility at equilibrium over 5000 experiments for each scenario. Except when rz is constructed using the “random noise” or “plateau” procedure and the magnitude of efficiency effects is small (asterisks), single mating tends to promote average worker sterility at equilibrium over double mating (the 0/0 denotes no worker sterility under either single or double mating). This can happen even if sterility is more likely to invade under double mating (for example, compare results of procedures i-iii in panel **(b)** versus panel **(c)**). Arrowheads beneath the x-axis show where parameters coincide with those used in panel **(b)**. The “magnitude of colony efficiency effects” is the standard deviation of normally-distributed variates used for constructing r_z_. For panels **(b)** and **(c)**, we assume *p_z_* = 0.2 + 0.8z. See Methods for details.

**Figure 3:**
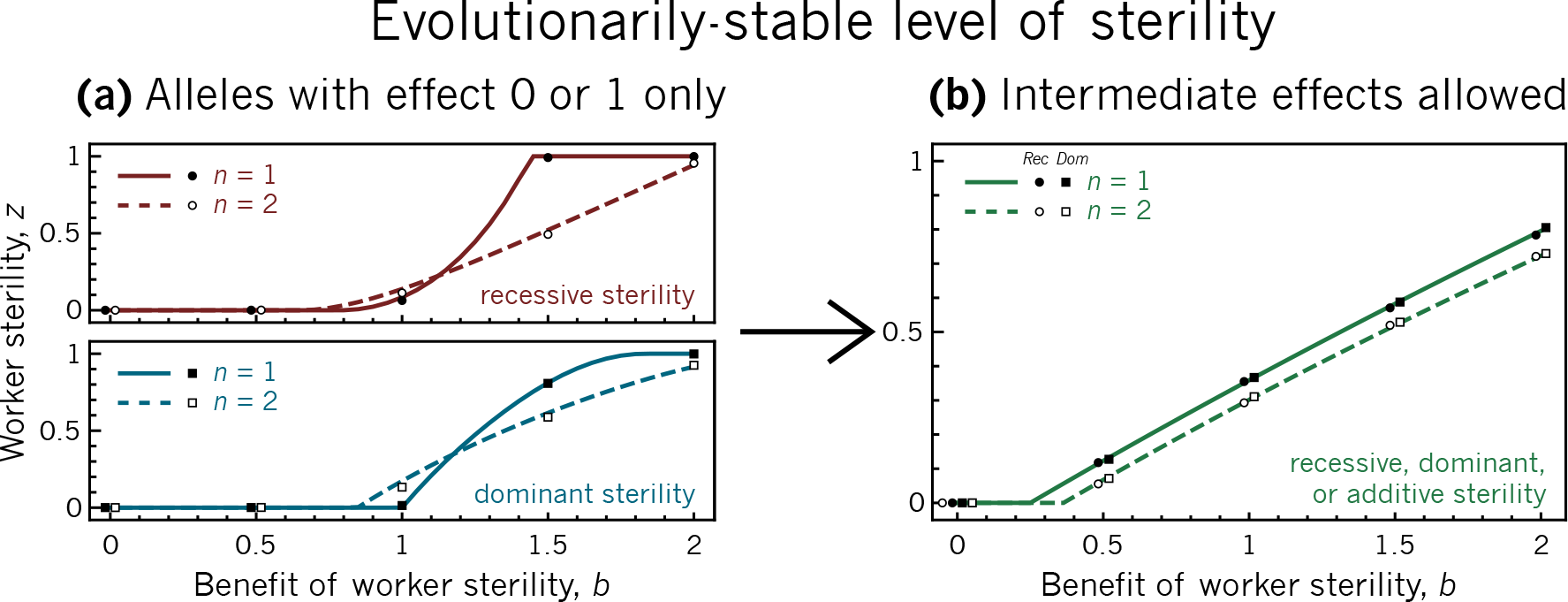
The evolutionarily-stable level of sterility under single versus double mating, for **(a)** constrained allelic variation, with recessive (top) versus dominant (bottom) sterility and **(b)** unconstrained allelic variation, regardless of whether sterility is recessive, dominant, or additive. **(a)** When allelic variation is constrained, double mating (dashed lines) can sometimes promote sterility over single mating (solid lines). But **(b)** when allelic variation is unconstrained, single mating always promotes sterility. Overlaid markers show results of a stochastic individual-based model (see Methods), matching well with the predicted evolutionarily-stable levels of worker sterility. To illustrate a scenario where constraints on heritable variation may lead to promiscuity promoting worker sterility over monogamy, we use the colony efficiency function *r_z_ =*1 + *bz* − *z*^2^, with a “benefit of worker sterility” term *bz* and a “decelerating” term −z^2^. For the proportion of male eggs laid by the queen, we again use *p_z_* =0.2 + 0.8z.

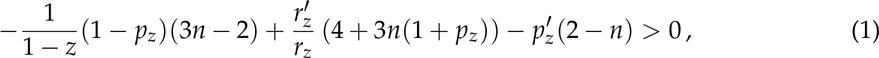

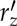 and 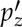 are the slopes of the *r*_z_ and *p*_z_ functions at z, respectively. Remarkably, this exact condition holds for both recessive and dominant genetics. Using this condition and a global stability analysis, we find that the ESS for helpful sterility is always highest under single mating—that is, over long-term evolution, monogamy always promotes helpful worker sterility (Fig. 3; see Methods).

Intuition for this exact population-genetics result may be obtained by recasting condition 1 in terms of inclusive fitness (Hamilton 1964). Accordingly, natural selection favours an increase to average sterility, z, when

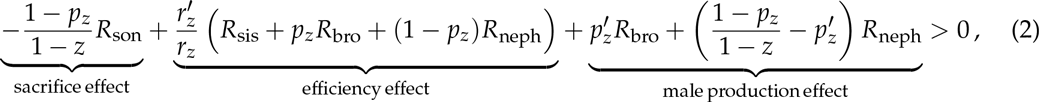

where 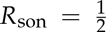,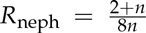, 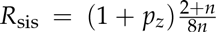, and 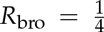 are the life-for-life relatedness of a worker to her son, her nephew (a random worker′s son), her reproductive sister, and her brother, respectively (Hamilton 1972). Note that promiscuity decreases worker relatedness to sisters and nephews, but not to sons or brothers. The left-hand side of condition 2 can be interpreted as the inclusive-fitness effect experienced by a focal worker who stops laying male eggs. The “sacrifice effect” captures the direct cost of her sterility, in that she forfeits her relative share 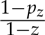 of all worker-laid males. The “efficiency effect” captures her impact on colony efficiency, which increases by a relative amount 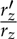 augmenting the production of her sisters and of colony-produced males, a proportion *p*_z_ of whom are her brothers, and a proportion 1 –*p*_z_ of whom are her nephews. And the “male production effect” captures her impact on the proportion of male eggs produced by the queen versus workers: her relative gain of brothers is 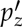, while her relative gain or loss of nephews exactly balances her forfeited sons and gained brothers.

Condition 2 clarifies the impact of monogamy upon helpful worker sterility: by increasing a worker′s relatedness to her nephews and sisters, monogamy increases her inclusive-fitness benefit of promoting colony efficiency, and by increasing a worker′s relatedness to her nephews, it increases her inclusive-fitness benefit of augmenting her fellow workers′ production of sons. Hence, overall, monogamy promotes helpful worker sterility. Condition 2 also clarifies how Olejarz *et al.’s* (2015) model differs from Boomsma(2007, 2009,2013) model for the evolution of eusociality: in Boomsma′s model, females trade away offspring for siblings as dispersers evolve into a non-totipotent worker caste, while in *Olejarz et al.′s* model, an existing non-totipotent worker caste trades away sons for brothers and nephews. Conditions 1 and 2 are exactly equivalent, are valid for recessive, dominant, or additive genetics, and can be obtained using standard kin-selection methodology (see Methods).

## Alternative ecological scenarios: monogamy promotes worker sterility

Finally, we consider some alternative scenarios for the evolution of worker non-reproduction, using a demographically-explicit model of queen-worker competition over egg-laying. Whether we investigate sex-blind egg replacement by workers, soldier sterility in claustral inbreeders, or the evolution of eusociality via female non-dispersal, we find that monogamy always promotes helpful worker sterility (Fig. 4). This conclusion also holds if we alternatively consider a diploid mode of inheritance (see Methods).

**Figure 4:**
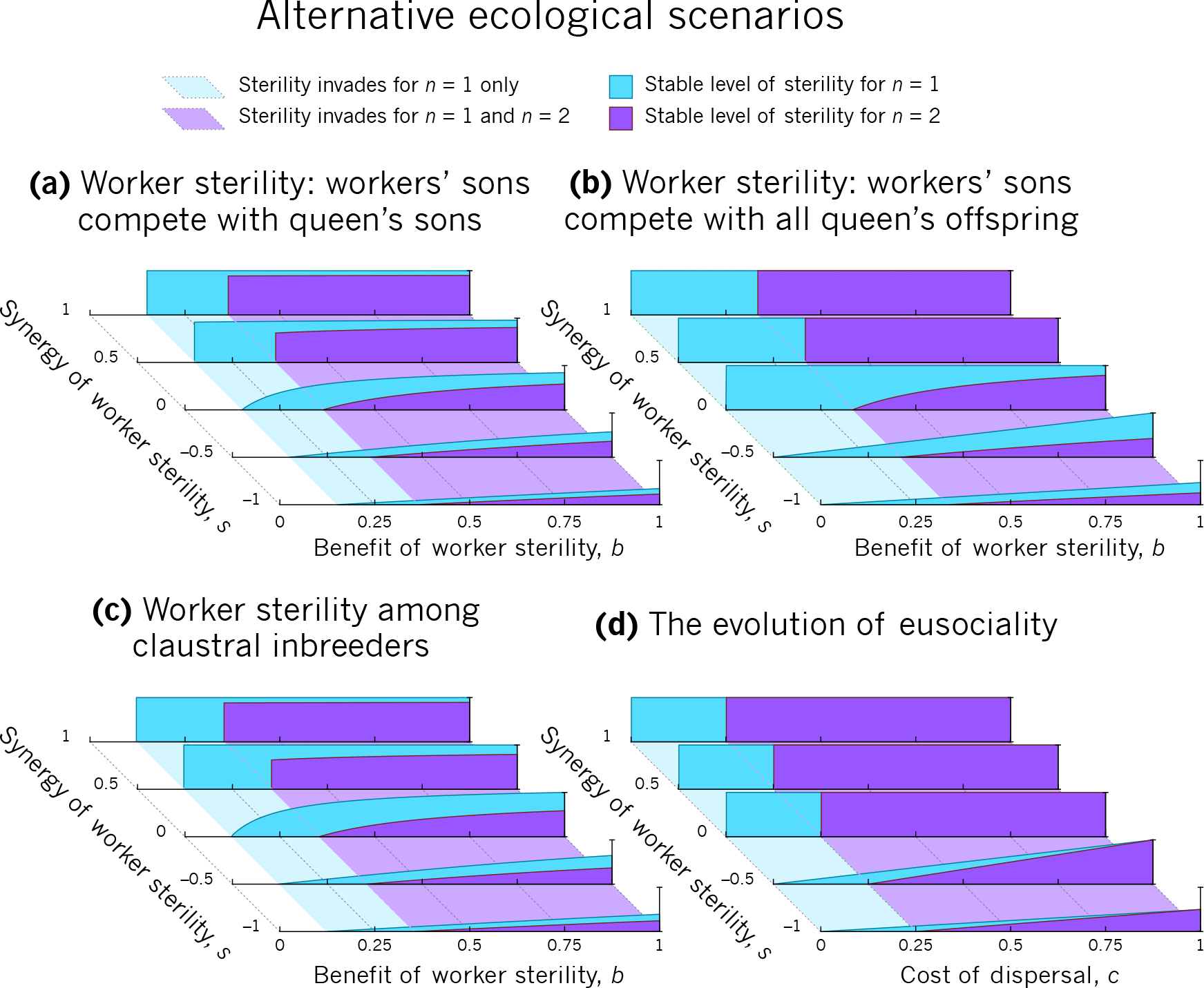
The evolution of worker sterility under alternative ecological scenarios. Here, we determine the stable level of worker sterility under four demographically-explicit models of worker sterility; see Methods for full details. **(a)** One possible assumption is that worker-laid males only compete with the queen′s sons (*cf.* Olejarz *et al.* 2015). In this case, monogamy promotes worker sterility over promiscuity. **(b)** It is also possible to assume that worker-laid males compete with the queen′s offspring of both sexes, and not just with the queen′s sons. In this case, monogamy promotes worker sterility over promiscuity. **(c)** In the gall-forming thrips, the foundress produces an initial brood of female and male soldiers, who may produce part of the next brood by inbreeding amongst themselves (Chapman *et al.* 2002). Female soldiers can sacrifice part of their reproductive potential to invest more in defending their nestmates. In this case, monogamy promotes worker sterility over promiscuity. **(d)** A possible model for the evolution of eusociality involves dispersing, fully-reproductive females evolving into sterile workers, who stay in the nest to help, producing no offspring (Boomsma 2007, 2009, 2013). In this case, monogamy promotes worker sterility over promiscuity. We show results for *k* =4 in **(a)** and *k* =2 in **(b)** and **(c)** (see Methods for details).

## Conclusion: monogamy promotes worker sterility

In seeming contrast to the predictions of inclusive-fitness theory, Olejarz *et al.′s* (2015) exact population-genetics analysis could not identify a consistent effect of monogamy on the evolution of worker sterility. This surprising result, if robust, would have not only overturned a considerable theoretical consensus, but would also have left a number of empirically-described patterns bereft of a predictive, explanatory framework. Happily, we have shown that by relaxing constraints on genetic variation (Fig. 1), considering the consequences of invasion rather than just its occurrence (Fig. 2), describing long-term evolutionarily-stable states (Fig. 3), and exploring a wide range of ecological scenarios (Fig. 4), a clear sterility-promoting effect of monogamy consistently emerges. Moreover, we have shown that the long-term evolutionary outcome is readily described, conceptualised, and explained by standard inclusive-fitness theory. In sum, a more comprehensive analysis based on Olejarz *et al.’s* (2015) exact population-genetics approach supports inclusive-fitness theory and its prediction that monogamy promotes the evolution of worker sterility.

## Acknowledgements

We thank Kevin Foster for helpful comments.Alternative ecological scenarios

## Methods

### Helpful versus harmful worker sterility and policing

Throughout the main text, our focus is on helpful worker sterility, where giving up some or all of her reproductive potential allows a worker to provide more help within her colony, as this biological assumption underpins most work on altruistic sib-rearing in social insects. However, the model of Olejarz *et al.* (2015), despite making strong genetic assumptions, makes few ecological assumptions about worker sterility, which means it may also describe harmful worker sterility. If worker sterility is harmful′namely, if worker sterility reduces colony efficiency and/or reduces other workers′ personal fitness—monogamy may inhibit worker sterility, depending on the overall impact of sterility on a worker′s inclusive fitness.

In this model, harmful worker sterility may occur via two routes—one operating through colony efficiency, *r_z_,* and one operating through the queen’s production of males, *p_z_.* The first case occurs when an increase in average worker sterility decreases colony efficiency—for example, if the sterility allele has a pleiotropic effect on worker condition which results in less-efficient work. In such a case, monogamy will inhibit the evolution of worker sterility relative to promiscuity, since promiscuity decreases relatedness between relatives, thereby lessening the harmful impact of sterility upon a worker′s inclusive fitness via colony efficiency.

The second case occurs when an an increase in a focal worker′s sterility harms the reproductive success of other workers. In the main text, we assume that when a worker becomes sterile, her forfeited sons are replaced partly by the queen′s sons and partly by her sisters′ sons, such that by forfeiting sons she gains both nephews and brothers. But if, due to the shape of the *p_z_* function, the queen gains a larger proportion of sons than the worker forfeits (that is, when 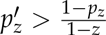),this “outsized gain” by the queen must be balanced by *decreased* male production by other workers, such that, by becoming sterile, the focal worker loses nephews overall. If the focal worker loses nephews by becoming sterile *(i.e.,* when 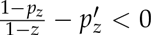; see condition 2), then promiscuity, by decreasing the worker′s relatedness to nephews, may promote this harmful form of worker sterility over monogamy, unless this relative cost of sterility is countered by a colony efficiency benefit of sterility, which would be largest in magnitude under monogamy.

This second form of harmful worker sterility is connected with worker policing—that is, when workers invest resources in preventing other workers from laying eggs (Ratnieks 1988; Ratnieks & Visscher 1989). If worker sterility is harmful, then a worker gives up part of her personal fitness in order to decrease the reproduction of her fellow workers; this is analogous to costly worker policing. Standard inclusive-fitness theory (Ratnieks 1988; Ratnieks & Visscher 1989; Ratnieks *et al.* 2006) and empirical evidence (Wenseleers & Ratnieks 2006a, 2006b) have emphasised that promiscuity promotes worker policing, so the result that this harmful form of worker sterility may be promoted by promiscuity is not at all surprising.

For non-incremental increases in sterility, the condition for harmful sterility becomes 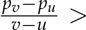 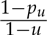 where *u* is the level of worker sterility in the monomorphic population before the mutant allele is introduced, and *v* is the level of worker sterility encoded by the mutant allele.

### Explicit population-genetics analysis

In Appendix A, we extend the methods of Olejarz *et al.* (2015) to consider the invasion of an allele with an arbitrary effect on worker sterility; the results of this analysis are presented here. We find that a recessive allele encoding worker sterility *v* can invade a population monomorphic for sterility *u* when

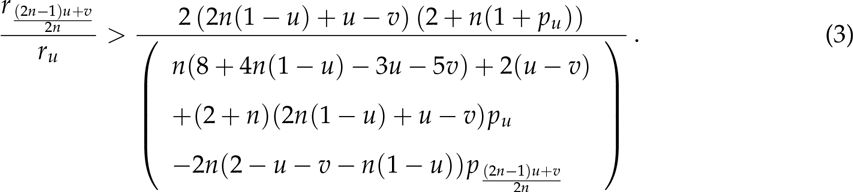

Similarly, we find that a dominant allele encoding worker sterility *v* can invade a population monomorphic for sterility *u* when

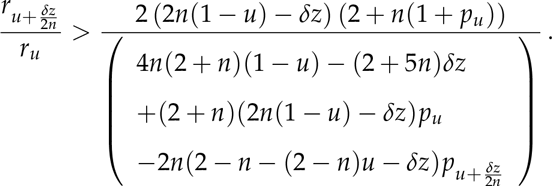

Note that conditions 3 and 4 give both the invasion and stability of a given level of sterility: that is, if a sterility allele with effect *v* can invade a population monomorphic for sterility u, then this is the same as saying that a population monomorphic for sterility *u* is not stable to invasion by a sterility allele with effect *v*. For example, substituting *n* = 1, *u* = 0, *v* = 1 into condition 3 yields the condition for the invasion of a recessive sterility allele under single mating from Olejarz *et al.* (2015; their condition 1), while substituting *n* = 1, *u* = 1, *v* = 0 into condition 4 yields the condition for the stability of a recessive sterility allele under single mating from Olejarz *et al.* (2015; their condition 3).

In order to find when natural selection will favour a small increase in sterility Sz, we make the substitution *v* = *u* + δ*z* into conditions 3 and 4 above. Then, by linearizing *r_z_* and *p_z_* around the point *z = u*, we can recast these conditions in terms of the value and slope of *r_z_* and *p_z_* at this point. More specifically, for a recessive sterility allele, substituting *v = u + δz* into condition 3 yields

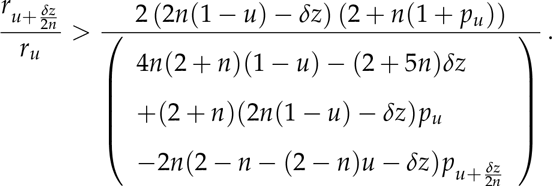

Linearizing *r_z_* and *p_z_* around z = *u*, we replace 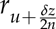 with 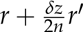, where r = *r*_u_ and 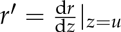.

Similarly, we replace 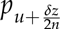 with 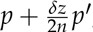, where *p = p_u_* and 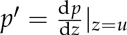. This yields

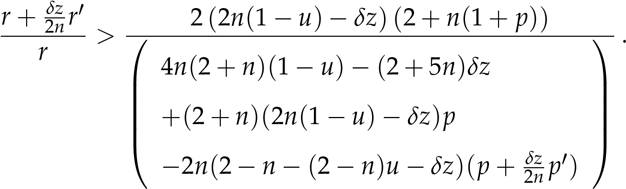

Eliminating the fractions on both sides, discarding terms of order *δz^2^* or higher, substituting *z* for *u* and simplifying yields 
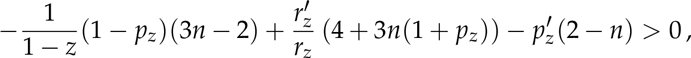
which is condition 1 of the main text.

Similarly, for a dominant sterility allele, substituting *v* = *u* + *δz* into condition 4 yields

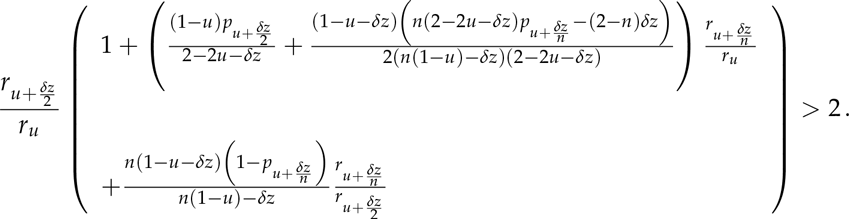

By linearizing r_z_ and *p*_z_ around z = u as above, we obtain 
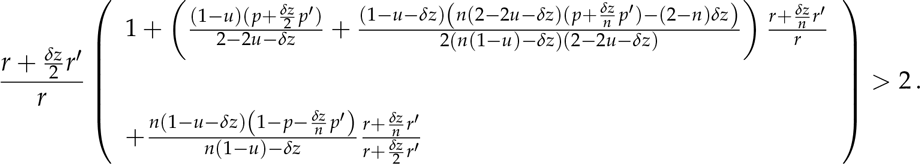

Expanding all terms, discarding terms of order δz^2^ or higher, substituting *z* for *u* and simplifying yields 
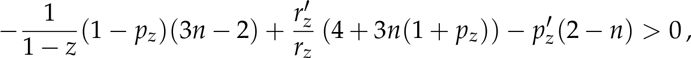
which, again, is condition 1 of the main text.

### Invasion of a dominant sterility allele

In the main text, we discuss why promiscuity can sometimes favour the invasion of a recessive worker sterility allele. The invasion of a dominant worker sterility allele is similar, but in this case there are two “mutant” mating types which determine whether sterility can invade: a heterozygous mutant female mating with *n* wild-type males, and a wild-type female mating with one mutant male and *n* — 1 wild-type males. These mating types produce colonies with a proportion 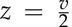 and 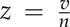 of sterile workers, respectively. Hence, for a sterility allele with full penetrance, under single mating (*n* = 1), it is the relative success of colonies with 50% and 100% sterile workers which determines whether a sterility allele with full penetrance can invade, while under double mating (*n* =2), only the relative success of colonies with 50% sterile workers determines whether a sterility allele with full penetrance can invade. Therefore, if the relative success of colonies with 100% sterile workers is low, this could be enough to disfavour the invasion of a fully-penetrant worker sterility allele into a wild-type population under single but not double mating. Nonetheless, for the scenario investigated by Olejarz *et al.* (2015, their Fig. 8), we find that single mating always promotes the invasion of dominant sterility over double mating (Fig. 5). Moreover, allowing for helpful-sterility alleles showing the full range of degrees of penetrance and domi-nance/recessivity, there is no scenario under which *some* sterility alleles can invade under double mating, and yet *no* sterility allele can invade under single mating.

### Numerical experiments

Olejarz *et al.* (2015) performed numerical experiments to see whether sterility was more likely to invade under single mating or double mating. To do so, they constructed randomly-generated *r_z_* functions according to one of two procedures. Here, we add to these procedures, bringing the number of possible methods for constructing the *r_z_* function to five (Fig. 2a). Each involves drawing four random variates—here, notated as *a, b, c*, and *d*—from a normal distribution with mean 0 and standard deviation *σ*. In all cases, we assume r_o_ = 1, and use the random variates to generate *r*_1/4_, *r*_1/2_, *r*_3/4_, and *r*_1_, which suffice to numerically integrate the evolutionary dynamics of worker sterility using the system of ODEs described by Olejarz *et al.* (2015). We restrict our attention here to the invasion of an allele encoding full sterility in its carriers, under either recessive or dominant genetics.

**Figure 5:**
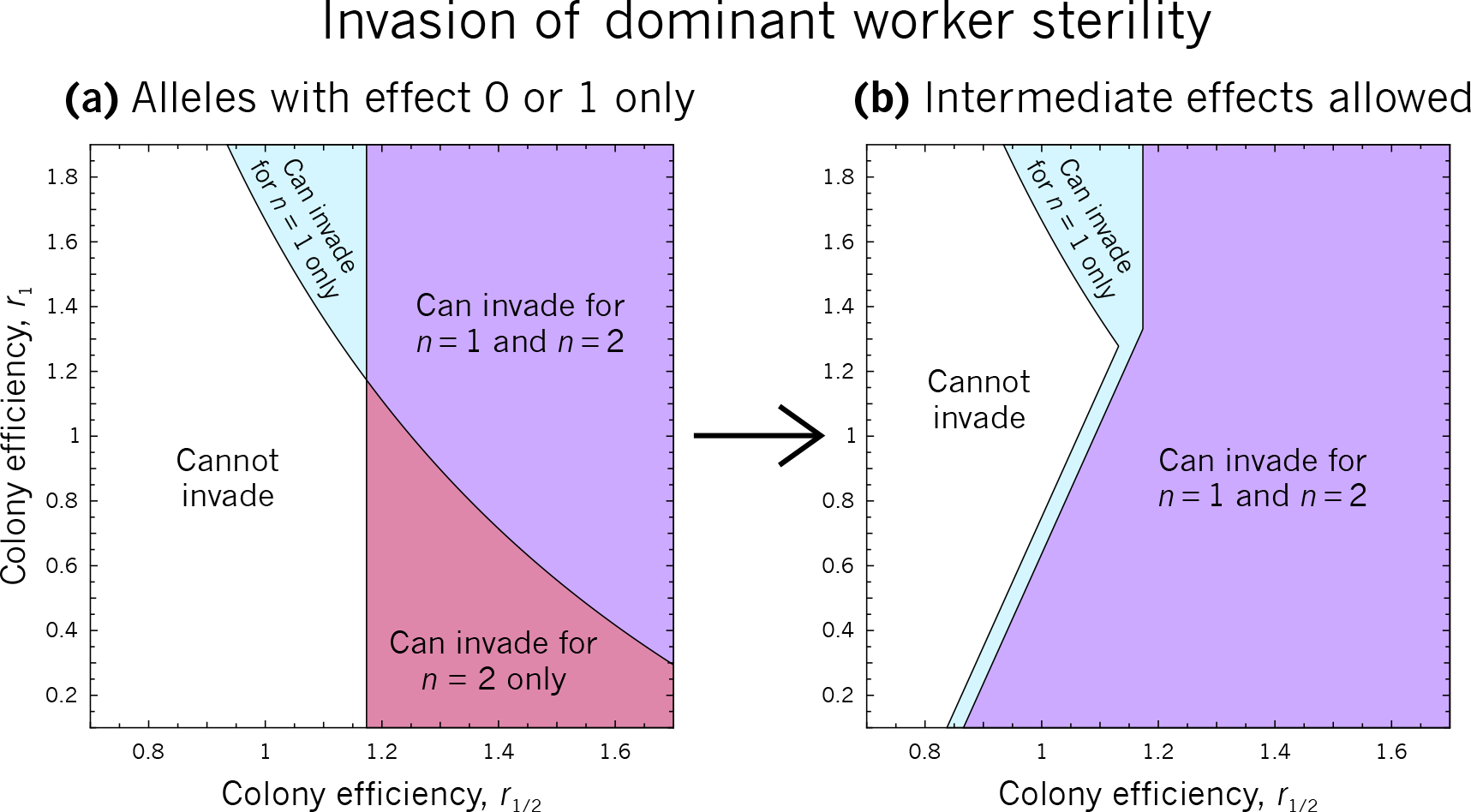
The invasion of worker sterility under dominant genetics, exploring the regions of parameter space where sterility can invade under single mating only, double mating only, both, or neither. **(a)** If we assume that only full-sterility alleles can arise, double mating sometimes promotes the invasion of sterility over single mating. But **(b)** if we assume that alleles encoding intermediate worker sterility can arise, double mating never promotes the invasion of sterility over single mating. In order to facilitate comparison with Figure 8 of Olejarz *et al.* (2015), we assume *p_z_* = 0.2 + 0.8z, and for *r_z_* we use the unique quadratic curve passing through the points specified by r_o_ = 1, r_1_/2, and r_1_.

The first procedure, “random noise”, is equivalent to Procedure 1 in Olejarz *et al.* (2015). Here, we set *r*_1/4_ = *r*_0_ + *a, r_1/2_ = *r*_0_* + b, *r*_3/4_ = *r*_0_ + c, and *r_1_ = *r*_0_* + d. Note that the four values are completely uncorrelated with each other; sequential values of r_z_ are independent from previous values, which is why we have named this procedure “random noise”. This procedure might generate plausible r_z_ functions for a population where every colony-level increase in worker sterility were to completely erase the effect of any previous increase in worker sterility, replacing it with a new, random effect.

The second procedure, “plateau”, is equivalent to Procedure 2 in Olejarz *et al.* (2015). Here, the values *r*_1/4_, *r*_1/2_, *r*_3/4_, and *r*_1_ are drawn from a correlated multivariate normal distribution. This can be simulated by transforming four uncorrelated normal variates; one way of doing this is by using the matrix 
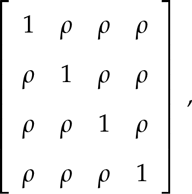
where *ρ* is the desired correlation between each variate. By multiplying the vector of uncorrelated variates by the Cholesky decomposition of this matrix, one obtains four correlated variates 
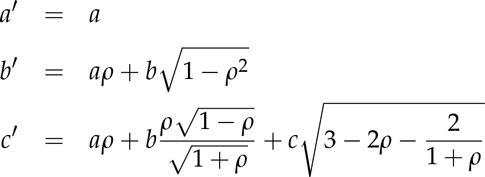

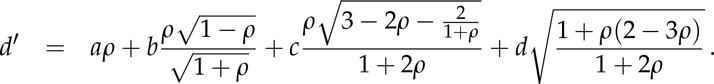

Now, we set r_1/4_ = *r*_0_ + *a′, r*_1/2_ = *r*_0_ + b′, *r*_3/4_ = *r*_0_ + c′, and *r*_1_ = *r*_0_ + d′. Note that, because the variables are correlated, the first “step” (from *r*_o_ to *r*_1/4_) tends to be larger in magnitude than subsequent “steps” (i.e., from *r*_1/4_ to *r*_1/2_, *r*_1/2_ to *r*_3/4_, or *r*_3/4_ to *r*_1_), which is why we have named this procedure “plateau”. This procedure might generate plausible *r*_z_ functions for a population in which worker sterility brings diminishing returns to colony productivity, where these diminishing returns happen to set in near z = 1/4.

Note that both the “random noise” and “plateau” procedures tend to produce *r_z_* functions that disadvantage single mating relative to double mating. For the “random noise” procedure, this is because although the procedure is just as likely to produce a peak at z = 1/2 (which would favour single mating) as at z = 1/4 (which would favour double mating), workers at z = 1/2 are typically “trading away” more male production than workers at z = ^1^/4 (since *p*_1/2_ > *p*_1/4_), yet, on average, they are receiving the same expected increase in productivity; hence, single mating is relatively disfavoured. And since the “plateau” procedure tends to produce colony efficiency functions with diminishing returns on worker sterility for colonies with z > 1/4, it is much more likely to produce an *r*_z_ function with a relative peak at z = 1/4 rather than a relative peak at z = 1/2.

The third procedure, “random steps”, sets each point in *r*_z_ to the value of the previous point plus a random perturbation: *r*_1/4_ = *r*_0_ + *a, r*_1/2_ = *r*_1/4_ + b, *r*_3/4_ = *r*_1/2_ + c, and *r*_1_ = *r*_3/4_ + d. This procedure might generate plausible *r*_z_ functions if each increase in worker sterility had a random increasing or decreasing effect on colony productivity. The fourth procedure, “increasing steps”, is similar, except steps are constrained to be positive: *r*_1/4_ = *r*_0_ + |a|, *r*_1/2_ = *r*_1/4_ + |b|, *r*_3/4_ = *r*_1/2_ + |c|, and r_1_ = *r*_3/4_ + |d|. This procedure might generate plausible *r*_z_ functions if each increase in worker sterility added a random increase to colony productivity. The fifth procedure, “linear”, uses a single normal variate to establish a constant step size for *r*_z_: *r*_1/4_ = *r*_0_ + a, *r*_1/2_ = *r*_1/4_ + a, *r*_3/4_ = r_1/2_ + a, and *r*_1_ = *r*_3/4_ + a. This procedure might generate plausible *r_z_* functions if each increase in worker sterility had a consistent increasing or decreasing effect on colony productivity. For each of these new procedures, later points in *r_z_* depend on earlier points, but there is no tendency for “steps” between points in *r_z_* to change in average magnitude.

In Fig. 2, we test each of these 5 procedures to see whether single or double mating promotes the invasion (Fig. 2b) or equilibrium level of sterility (Fig. 2c) more, for recessive versus dominant sterility. The form of *p_z_* we use (*p_z_* = *k* + (1 — *k)z*, with *k* = 0.2), chosen for comparison with the numerical experiments of Olejarz *et al.* (2015, their (Table 1), prevents worker sterility from resulting in a net loss of nephews (see *Helpful versus harmful worker sterility and policing,* above). Beneath the bar charts in Fig. 2d, we show the percentage of experiments for which the exclusive invasion of sterility under either single or double mating occurred with an initially-decelerating *r_z_ (i.e.,* where *r*_1/2_ — *r*_1/4_ < *r*_1/4_ — r_0_). Note that, for these values of *p_z_,* double mating only promotes the invasion of sterility over single mating when *r_z_* is initially-decelerating. In Fig. 2c, error bars show bootstrapped 95% confidence intervals for average worker sterility.

### ESS analysis

By setting the left-hand side of condition (2) to zero, it is possible to find a convergence-stable point (Davies *et al.* 2016) for worker sterility. At these points, natural selection will not favour the invasion of an allele encoding either a small increase or a small decrease to worker sterility (*i.e.*, convergence-stable points are stable to small perturbations); moreover, for a population playing a strategy that is close to a convergence-stable point, natural selection will favour the invasion of strategies between the population strategy and the convergence-stable point (*i.e.*, convergence-stable states are reachable from nearby states). However, a convergence-stable point is only an evolutionarily-stable strategy (ESS) if *no* alternative allele can invade at this point. Therefore, in order to find a true ESS, we treat convergence-stable points as “candidate ESSs”, then use conditions 3 and 4 to determine whether any alternative allele can invade a population monomorphic for the candidate ESS under the appropriate regime of dominance or recessivity. If no alternative allele can invade, the candidate ESS is a true ESS. In Figure 3, true ESSs are shown.

Note that it is possible for an ESS to *not* be convergence-stable, and this method will not identify such states. However, we are only interested in ESSs that are reachable, *i.e.*, both convergence-stable and evolutionarily-stable. Such strategies are called “continuously-stable strategies” (CSSs; Eshel 1983).

### Demographically-explicit ecological scenarios

In Appendix B, we develop a general kin-selection model for the evolution of worker sterility. This analysis can be used to investigate a variety of ecological scenarios. Here, we present four such scenarios for the evolution of worker sterility.

#### Scenario A. Workers′ sons replace queen′s sons

In this scenario, we assume that non-sterile workers replace the queen′s sons with their own sons, as in the model of Olejarz *et al.* (2015). Following these assumptions, we find that natural selection will favour an increase to worker sterility, z, when 
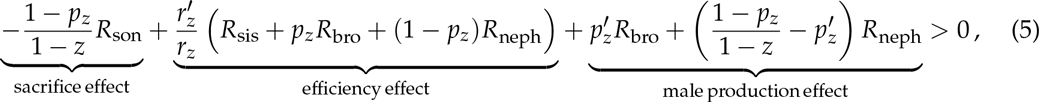
where 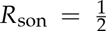, 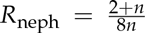, 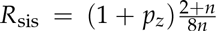, and 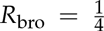. As explained in the main text, the left-hand side of condition 5 can be interpreted as the inclusive-fitness effect experienced by a worker who stops laying male eggs. The “sacrifice effect” captures the direct cost of her sterility, in that she forfeits her relative share 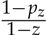 of all worker-laid males. The “efficiency effect” captures her impact on colony efficiency, which increases by a relative amount 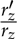, augmenting the production of her sisters and of colony-produced males, a proportion *P_z_* of whom are her brothers, and a proportion 1 —*P_z_* of whom are her nephews. And the “male production effect” captures her impact on the proportion of male eggs produced by the queen versus workers: her relative gain of brothers is 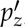, while her relative gain or loss of nephews exactly balances her forfeited sons and her gained brothers.

Similarly, natural selection favours an increase to the queen′s sex allocation, *x* (her proportion of resources allocated to daughters), when

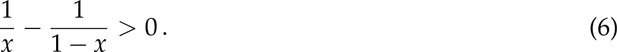

That is, natural selection favours an increased investment into daughters when x < 1/2, and a decreased investment into daughters when x > 1/2, such that an even sex ratio is favoured overall, regardless of worker sterility.

#### B. Workers′ sons compete with all queen′s offspring

It is also possible to assume that, rather than only displacing the queen′s sons, workers′ sons compete with the queen′s sons and daughters equally. This scenario may apply if workers do not discern between fertilised and unfertilised eggs when they replace the queen′s eggs with their own; alternatively, it may apply if rather than replacing the queen′s eggs, the workers simply lay their eggs in the communal nest, and all queen-produced and worker-produced offspring have the same expected survival. Following these assumptions, we find that natural selection will favour an increase to worker sterility, z, when 
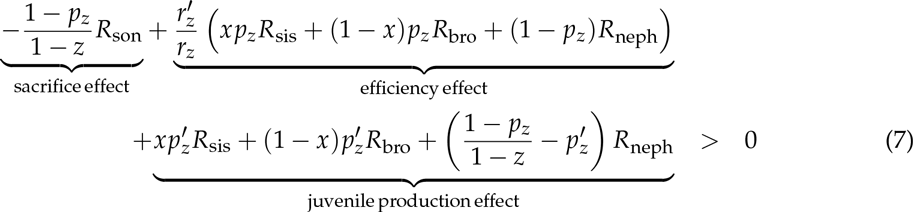
where *p*_z_ is the proportion of all juveniles on the patch that are produced by the queen, 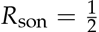, 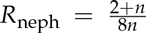, 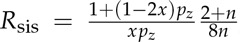, and 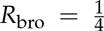. In this model, queen sex allocation alters the relative reproductive value of a female compared to that of a male, 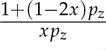 (the product of the relative reproductive value of all females compared to that of all males, 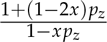, and the number of females relative to the number of males, 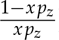), which comes into the expression for R_sis_. Similarly to condition 5, the left-hand side of condition 7 can be interpreted as the inclusivefitness effect experienced by a worker who stops laying male eggs. Here, the “sacrifice effect” captures the direct cost of her sterility, in that she forfeits her relative share 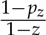 of all worker-laid males. The “efficiency effect” captures her impact on colony efficiency, which increases by a relative amount 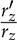, a proportion *xp*_z_ of which goes toward sisters, (1 — x)*p*_z_ toward brothers, and 1 —*p_z_* toward nephews. And the “juvenile production effect” captures her impact on the proportion of eggs produced by the queen versus workers: her relative gain of sisters is 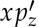, and her relative gain of brothers is 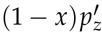, and so her relative gain of nephews exactly balances her lost sons, less her gained brothers and sisters.

In this scenario, queen sex allocation is not independent of worker sterility. We find that natural selection favours an increase to the queen’s investment in daughters, *x*, when 
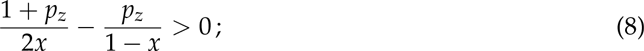
hence, when all colony offspring are queen-laid (*P_z_* = 1), the queen favours an even sex ratio (x = ^1^/2), but as the proportion of colony offspring laid by workers increases, the queen favours an increasingly female-biased sex ratio. Specifically, the equilibrium sex ratio is 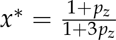.

#### Scenario C. Worker sterility among claustral inbreeders

Here, we assume that the queen produces a first brood of female and male soldiers, who mate amongst themselves; the second brood of female and male dispersers is partly produced by the queen and partly produced by the soldiers, as in the gall-forming social thrips (Chapman 2002). For simplicity, we assume here that queens and soldiers produce an even sex ratio for the second brood, but allowing sex ratio evolution does not change the results qualitatively (not shown). Following these assumptions, we find that natural selection favours an increase to the sterility of female soldiers, *z*, when 
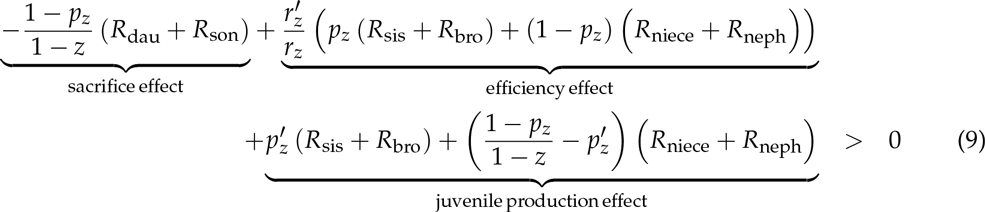
where, under haplodiploidy, 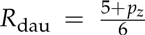, 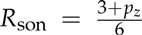, 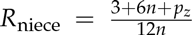, 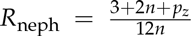, 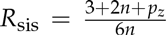, and 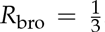. Because this scenario does not require arrhenotokous partheno-genesis of males, it also applies to diploid populations. Under diploidy, 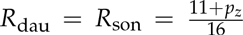 and 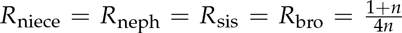 (Fig. 6a). Similarly to condition 7, the left-hand side of condition 9 can be interpreted as the inclusive-fitness effect experienced by a worker who stops 452 laying male eggs; but in condition 9, the female worker’s “sacrifice effect” involves giving up both daughters and sons; the “efficiency effect” involves an increase in both niece and nephew production as well as sister and brother production; and the “juvenile production effect” involves the focal worker gaining both sisters and brothers, while her gain or loss of nieces and nephews balances her forfeited offspring and her gained siblings.

**Figure 6:**
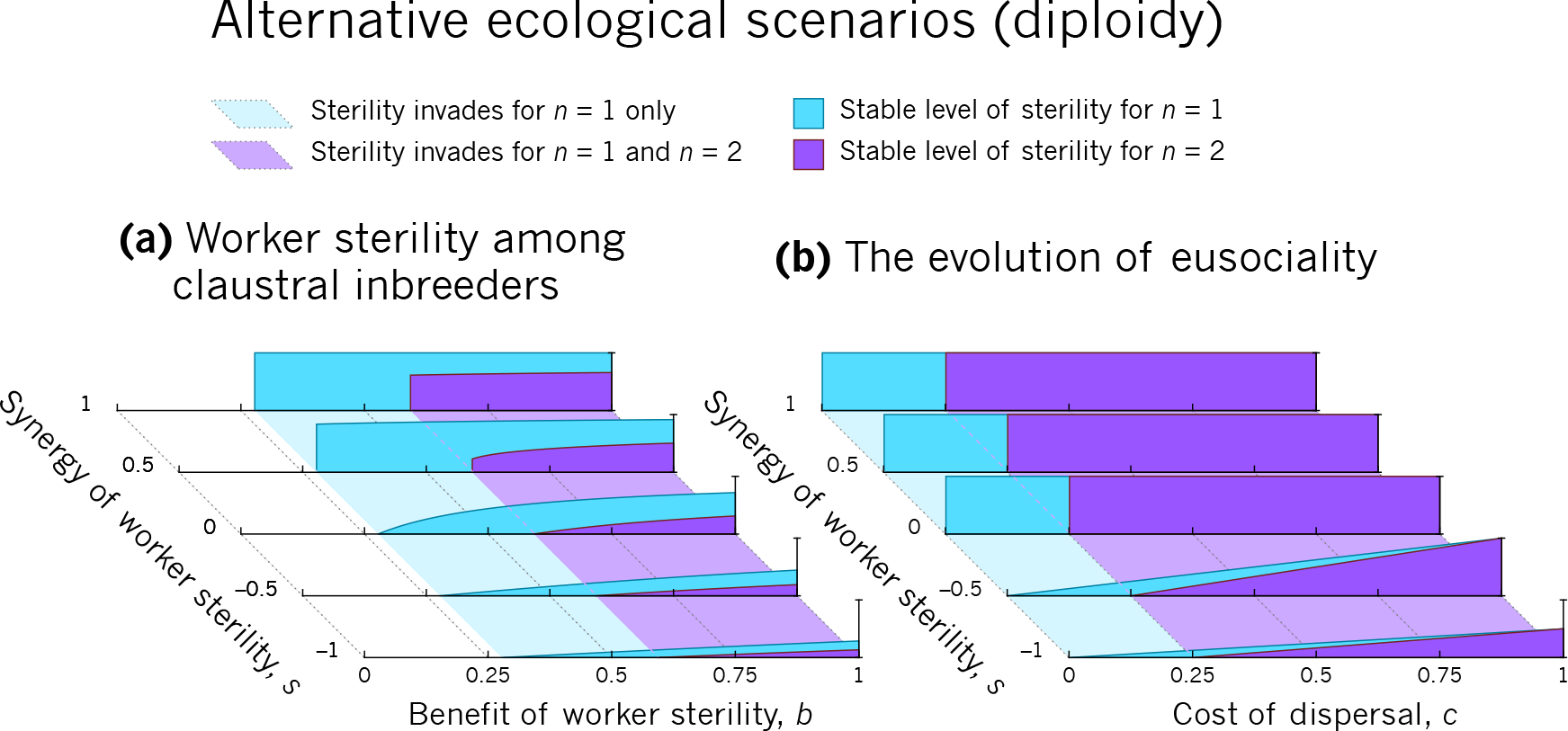
The evolution of worker sterility under alternative ecological scenarios, for diploidy. Here, we determine the stable level of worker sterility under two demographically-explicit models of worker sterility; see Methods for full details. **(a)** For claustral inbreeders under diploidy, monogamy promotes worker sterility over promiscuity; we show results for k = 4 here. **(b)** For the evolution of eusociality via non-dispersing female workers under diploidy, monogamy promotes worker sterility over promiscuity.

#### Scenario D. The evolution of eusociality

Here, we assume that the queen produces and provisions a first brood of females, and then produces a second batch of female and male eggs. Each first-brood female can either disperse—leave the nest, mate, and produce female and male offspring on her own—or work—stay in the nest and help to raise the queen’s second-brood offspring without producing any offspring of her own. We assume that each worker can raise *b* siblings, on average, in her natal nest, and that each disperser can raise *b*(1 — *c*) offspring, on average, in her newly-founded nest, where c represents the cost of dispersal; and, additionally, that workers may synergistically or antagonistically interact according to the parameter s, such that if the total number of female workers is Kz, then in total workers can raise *Kzb(1 + sz)* of the queen’s second-brood offspring. This model is similar to the one considered by Boomsma (2007, 2009, 2013) for the evolution of eusociality. Following these assumptions, we find that natural selection will favour an increase to worker sterility, *z*, when 
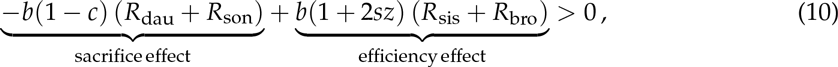
where 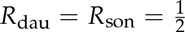, 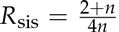, and 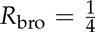. As with scenario C, this scenario also applies to diploid populations; under diploidy, 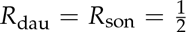 and 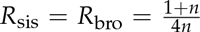 (Fig. 6b). When z = 0, this condition reduces to 
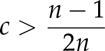
under both haplodiploidy and diploidy; that is, under strict monogamy (*n* = 1), any marginal benefit of rearing siblings over offspring (for example, any non-zero cost of dispersal, mating, or nest founding) suffices to favour the invasion of sterile workers, regardless of the level of worker synergy, s; but with any level of multiple mating (*n* > 1), a threshold dispersal cost of at least 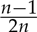 is required for natural selection to favour the invasion of sterile workers (Fig. 4d; Fig. 6b). In other words, only marginal efficiency gains are needed for worker sterility to invade under strict monogamy (Boomsma 2007, 2009, 2013).

#### Explicit forms for *r*_z_ and *p*_z_

Scenarios A, B, and C above are independent of the particular r_z_ and p_z_ functions used. However, for preparing Figs. 4 and 6, we used the explicit forms 
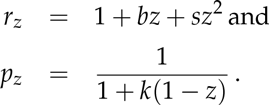

The *r*_z_ function above has three components: a baseline efficiency of 1; bz, representing a linear fitness benefit for each sterile worker; and sz^2^, representing an “interaction effect” of worker sterility. We use the parameter s to examine scenarios where multiple sterile workers results in either synergy (*s* > 0) or diminishing returns (*s* < 0) to colony productivity.

The *p*_z_ function given above corresponds to a model in which the queen and *k*(1 — *z*) reproductive workers each take an equal share of offspring production. Alternatively, *k* can capture not only the total number of workers but also their ability to control offspring production relative to the queen; for example, halving k could represent either a halving in the number of workers or a halving of their relative ability to control offspring production, keeping the number of workers constant.

A function of this form can also model more complicated demographic processes: for example, if we assume that there are N workers, each of whom replaces a random egg with their own at rate W, while the queen can replace a workers’ egg with her own at rate *Q*, then the form above gives the proportion of eggs produced by the queen at equilibrium when 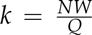. In models where worker-laid and queen-laid individuals compete equally, regardless of their sex, production of eggs and replacement of eggs will often be equivalent processes: that is, the form given above for *p*_z_ also holds if workers, rather than replacing the queen’s eggs, simply lay their own eggs in the communal nest without replacement. In that case, the *r*_z_ function would capture the overall production and survival of eggs.

#### Stable level of sterility

For Fig. 4, we determine the convergence-stable point (Davies et al. 2016) for sterility by numerically integrating the selection gradients for sterility and sex allocation (left-hand sides of conditions 5-10). First, we set the sex ratio to *x* = *x*̄ = 1/2 and allow it to evolve in the absence of worker sterility (*Z* = *z* = *z*̄ = 0) until it reaches its equilibrium value. Then, we allow both the sex ratio and sterility to coevolve, until equilibrium is reached for both traits.

### Stochastic individual-based model

To verify the results of our kin selection analysis (Fig. 3), we implemented a stochastic individual-based model in C++. Here, each individual comprises a locus encoding their breeding value for worker sterility, Z. The locus comprises one or two genes, depending on whether the individual is haploid or diploid, and each gene is represented by a real number γ∊ [0,1]. Breeding values are determined by averaging genic values: hence, a haploid individual with genotype 7 has breeding value Z = γ, while a diploid individual with genotype γ_1_,γ_2_ has breeding value Z = (γ_1_ + γ_2_)/2.

At the beginning of each generation, *M* mated females each produce *K* female workers on their home patch. Each worker has a probability Z of being sterile. The patch average sterility z determines the colony productivity *r*_z_ and the proportion of males produced by the queen *p*_z_. The next generation of breeders is then produced: first, a patch is randomly selected from the population with probability proportional to its colony efficiency, *r*_z_, and a female is produced by the queen on that patch; then, another n patches are randomly selected with replacement, with probability proportional to their colony efficiency, and each of these n patches produces a male (from the queen with probability *p*_z_, or from a random reproductive worker on that patch with probability 1 — *p*_z_); the female mates with these n males, and this process is performed M times, at which point all the M mated females replace the foundresses of existing patches. All other individuals on each patch die, returning the population to the beginning of the life cycle.

Simulations start with a monomorphic population in which all γ = 0, and hence Z = 0 for each individual. A gene in a newly-produced individual has a 1% probability of mutating, in which case its genic value changes from γ to γ′ = max(0,min(γ + δ,1)), where *δ* is drawn from a normal distribution with mean 0 and standard deviation 0.01. We validated this stochastic individual-based model by using it to verify the analytical conditions of Olejarz *et al.* (2015; not shown).

## Appendix A: Explicit population-genetics analysis

Here, we analyse the invasion of a sterility allele into a wild-type population. The population is initially monomorphic for an allele *A* encoding sterility with penetrance 0≤*u*≤1, and a rare mutant allele a is introduced which encodes sterility with penetrance 0≤*v*≤1. Throughout, we closely follow the approach of Olejarz *et al.* (2015), whose analysis is equivalent to ours with the assumptions that *u* and *v* are restricted to either 0 or 1.

We denote colony types by the genotype of the queen and the genotypes of her mating partners. Hence, X*_AA,m_* is the frequency of colonies with an AA queen, m mutant (a) males, and *n* — *m* wild-type (A) males; similarly for X*_Aa,m_* and X*_aa,m_*. At any given time step, we also keep track of the number of reproductive females of each genotype—x*_AA_*, *x_Aa_*, and *x_aa_*—and the number of reproductive males of each genotype—*y*_*A*_ and *y*_*a*_. Matings between reproductives lead to the establishment of new colonies; hence, the evolutionary dynamics of colony types are captured by: 
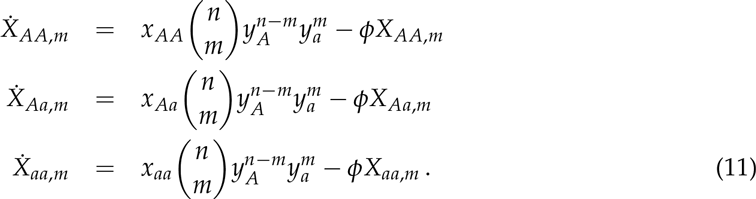

That is, the rate of establishment of new *AA*, *m* colonies is proportional to the frequency of reproductive *AA* females, multiplied by their probability of mating with exactly *n* — *m* wild-type males and *m* mutant males; similarly for Aa, *m* and *aa*, *m* colonies.

The death rate of existing colonies, ∅, is defined as 
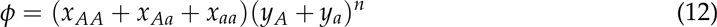
in order to enforce a density constraint, namely: 
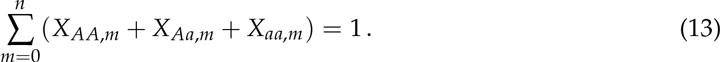

## Reproductives if the mutant allele is dominant

When the mutant allele is dominant, the production of each type of reproductive female (*x_AA_*, *x*_*Aa*_, *x_aa_*) and male (*y_A_*, *y_a_*) is: 
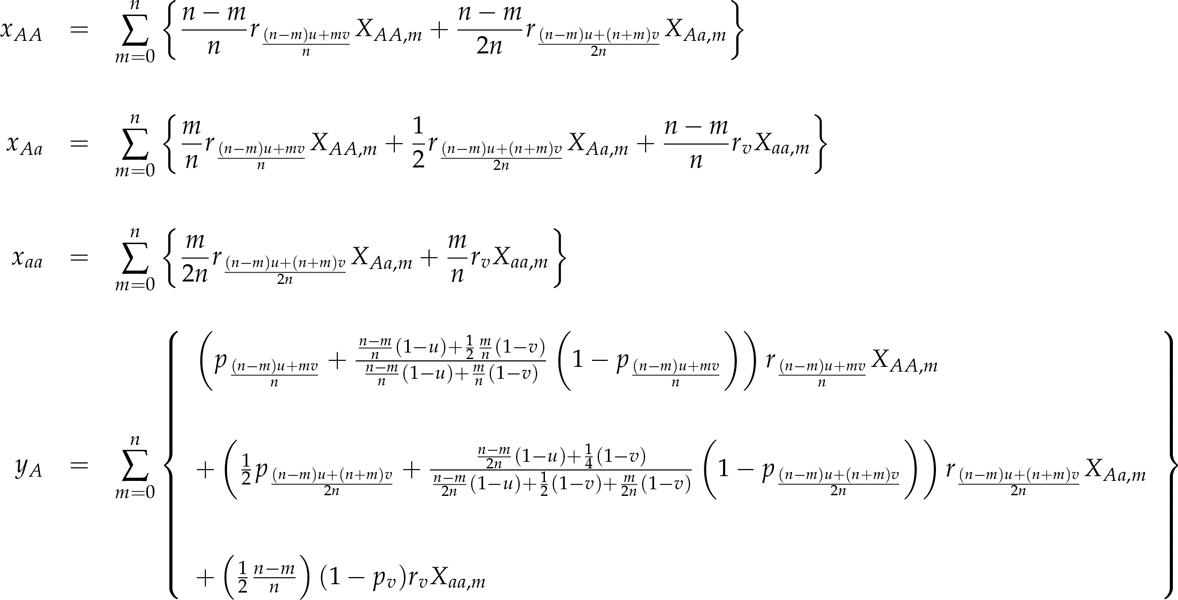

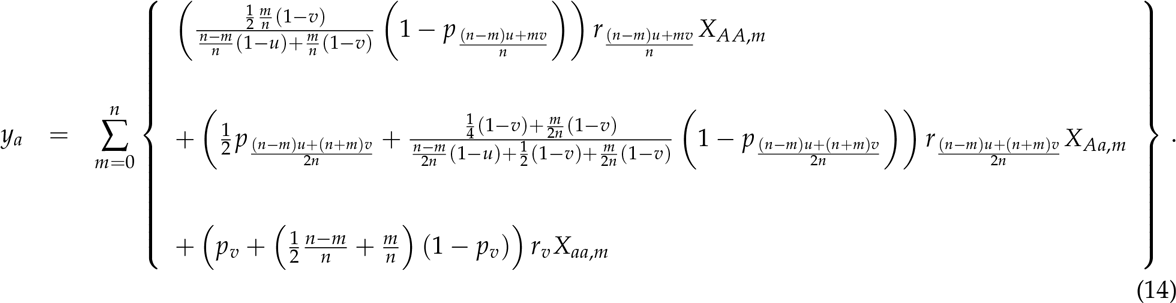

These equations can be understood as follows. First, note that in an *AA*, *m* colony, a fraction 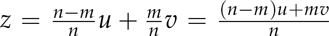 of workers will be sterile (*AA* workers with probability *u*, and *Aa* workers with probability *v*); in an *Aa*, *m* colony, a fraction z 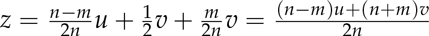 of workers will be sterile (*AA* workers with probability with probability *u*, and *Aa* and *aa* workers with probability *v*); and in an *aa*, *m* colony, a fraction 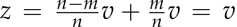 of workers will be sterile (*Aa* and *aa* workers with probability *v*). That is why these values of z as subscripts to the *r*_z_ and *p*_z_ functions are always associated, above, with their associated colony frequencies, *X_AA,m_*, *X_Aa,m_*, and *X_aa,m_*, respectively.

For female reproductives, each separate term within the curly braces above combines three elements; we will take the first term in curly braces in the *x_AA_* line, 
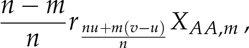
as an example. The three elements are the frequency of a given colony type (*i.e.*, *X_AA,m_*); the productivity of that colony type, as a function of the fraction of sterile workers within colonies of that type (*i.e.*, 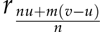); and the fraction of females and/or males produced by that colony type with the corresponding genotype (*i.e.*, a fraction 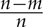 of females produced in *AA*, *m* colonies have genotype *AA*, which is why they add to the quantity *x_AA_*). Each term within equation 14 can be broken down in this way.

Accordingly, the production of female reproductives can be understood as follows: *AA*, *m* colonies produce 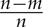 *AA* females and 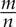 *Aa* females; *Aa*, *m* colonies produce 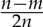 *AA* females, 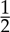 *Aa* females, and 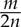 aa females; and *aa*, *m* colonies produce 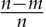Aa females and 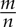 *aa* females.

Male production is more complicated, since both queens and workers produce males, but the principle is the same. We will take the first term in curly braces in the *y_A_* line, 
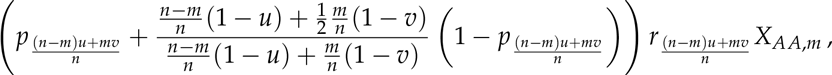
as an example. Here, the overall productivity of *AA*, *m* colonies (*i.e.*, 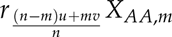) goes to-ward the production of both the queen’s sons and workers’ sons. In particular, the queen is *AA*, so all her sons have genotype *A*, and the queen produces a fraction *p* 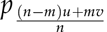 of males in the colony. Simultaneously, the workers—whose sons comprise a fraction 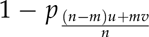 of colony male production—are 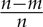 AA and 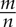 *Aa*; in the former group, workers are reproductive with probability 1 − *u*, while in the latter group, workers are reproductive with probability 1 − *v*; and all the sons of the first group will be *A*, while only half of the sons of the second group will be *A*. Hence, overall, a fraction 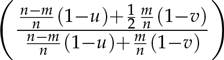 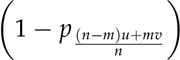 of males produced in *AA*, *m* colonies are *A* males produced by workers. Note that the expressions for *y*_A_ and *y*_a_ can be further simplified, but we have left them in the form above to maximise clarity.

Accordingly, the production of male reproductives can be understood as follows. In *AA*, *m* colonies, the queen’s sons are all *A*; all of the sons of *AA* workers and half of the sons of *Aa* workers are *A*, while the other half of the sons of *Aa* workers are a. In *Aa*, *m* colonies, the queen’s sons are half *A* and half a; all of the sons of *AA* workers and half of the sons of *Aa* workers are *A*, while the other half of the sons of *Aa* workers and all of the sons of *aa* workers are a. Finally, in *aa*, *m* colonies, the queen’s sons are all *a*; half of the sons of *Aa* workers are *A*, while the other half of the sons of *Aa* workers and all the sons of aa workers are *a*.

## Reproductives if the mutant allele is recessive population-genetics analysis

Along similar principles, when the mutant allele is recessive, the production of each type of reproductive female and male is: 
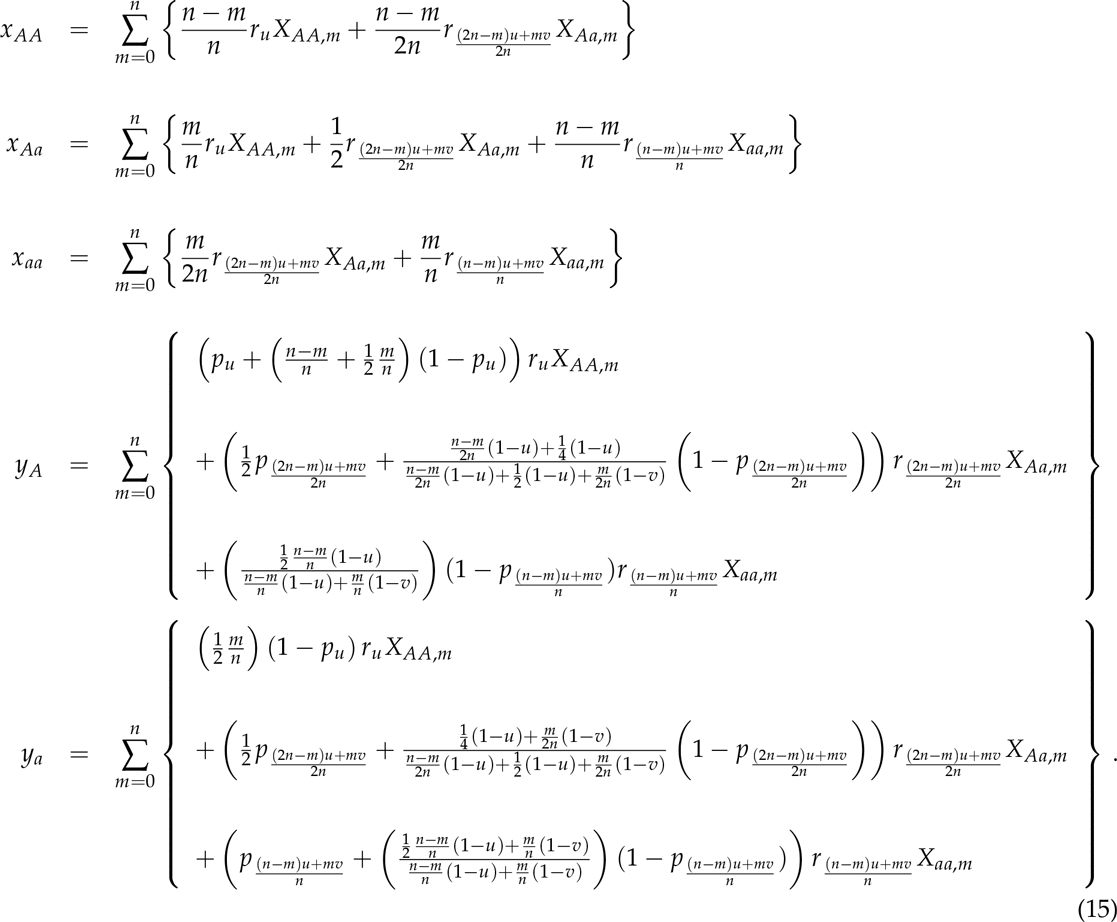

These equations can be understood similarly to equation 14; in fact, they are identical, except for two general changes. First, the subscripts to *r*_z_ and *p*_z_ are different, because the mutant allele is recessive instead of dominant, which results in different proportions of sterile workers in colonies of each type: in an *AA*, *m* colony, a fraction 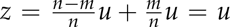 of workers will be sterile; in an *Aa*, *m* colony, a fraction 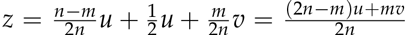 of workers will be sterile; and in an aa, *m* colony, a fraction 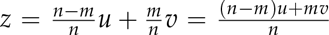 of workers will be sterile. Second, because of these differing proportions of sterile workers, the production of sons by workers is different, so the coefficients of 1 − *p_z_* in the fourth and fifth lines are different.

## Condition for invasion of a dominant mutant sterility allele

Continuing to follow the approach of Olejarz et al. (2015): for a dominant mutant sterility allele, whether the allele increases in frequency from rarity is governed by the behaviour of *AA*, 0, *AA*, 1, and *Aa*, 0 colonies. Colony types with more copies of the mutant allele are rarer, and so will have a negligible effect on invasion. Therefore, from equation 11, we need only consider: 
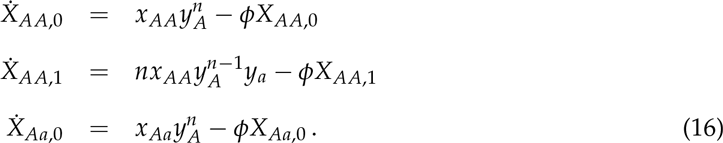

We start with a wild-type population (*X_AA_*,_0_ = 1) and introduce a small perturbation of magni-tude *ɛ* ≪ 1. Considering the density constraint (equation 13), and only keeping terms up to order *ɛ*, this gives 
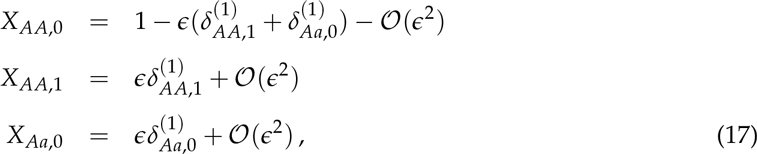
which implies that 
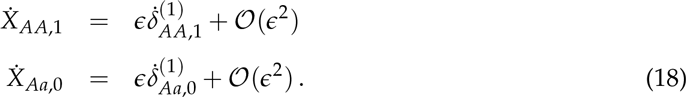

Substituting 17 into 14, and keeping terms only up to order ɛ, gives 
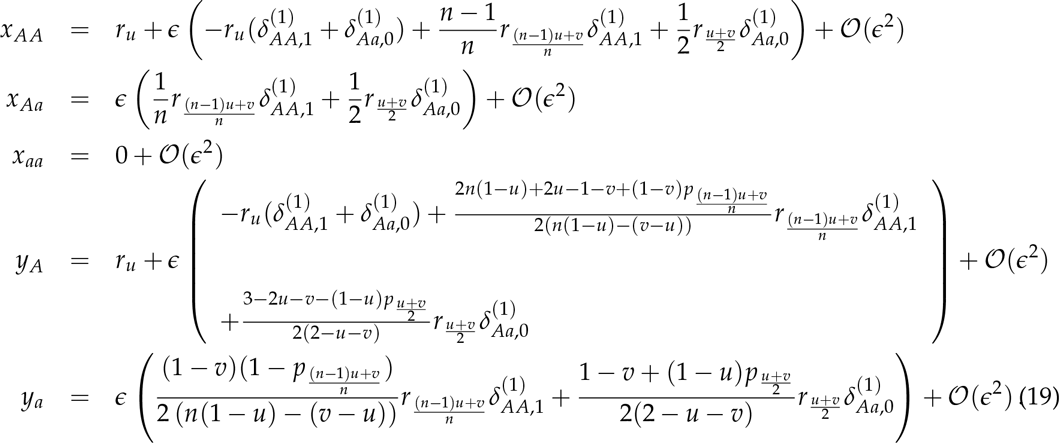

Finally, substituting 12,18, and 19 into 16 and discarding powers of ɛ^2^ or higher gives 
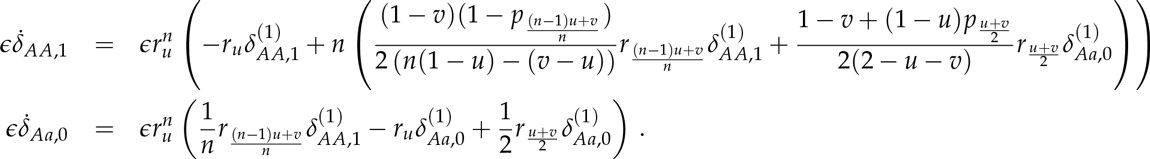

This can be rewritten in matrix form as 
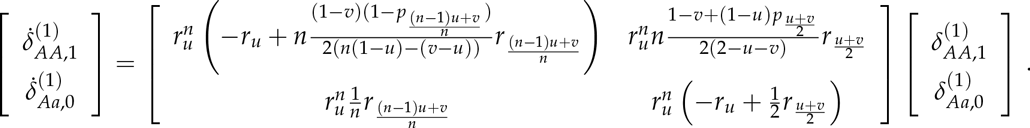

If the dominant eigenvalue of the above matrix is greater than zero, then a dominant sterility allele with penetrance *v* can invade a population monomorphic for sterility with penetrance *u*.

This condition, after simplification, is 
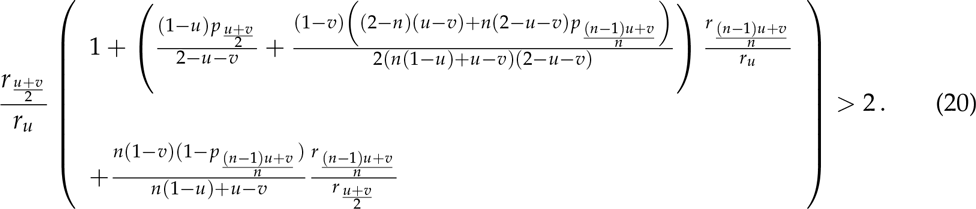

## Condition for invasion of a recessive mutant sterility allele

For a recessive mutant sterility allele, whether the allele increases in frequency from rarity is governed by the behaviour of *AA*, 0, *AA*, 1, *Aa*, 0, *AA*, 2, *Aa*, 1, and *aa*, 0 colonies. Colony types with more copies of the mutant allele are rarer, and so will have a negligible effect on invasion. Therefore, from equation 11, we need only consider: 
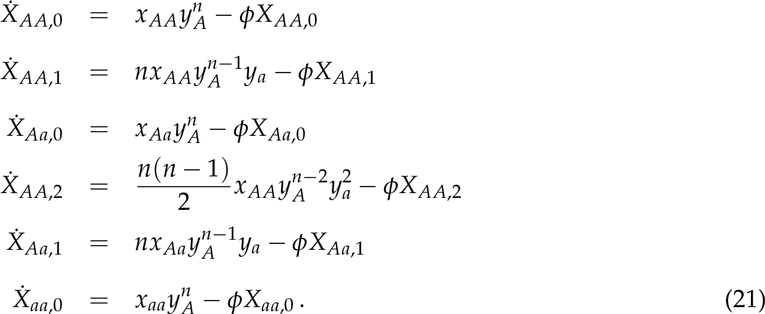

We start with a wild-type population (*X_AA,0_* = 1) and introduce a small perturbation of mag-nitude ɛ ≪ 1. Considering the density constraint (equation 13), and only keeping terms up to order ɛ^2^ (since terms of order ɛ alone are not sufficient to determine whether the recessive allele invades), this gives 
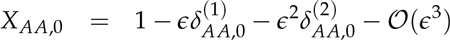

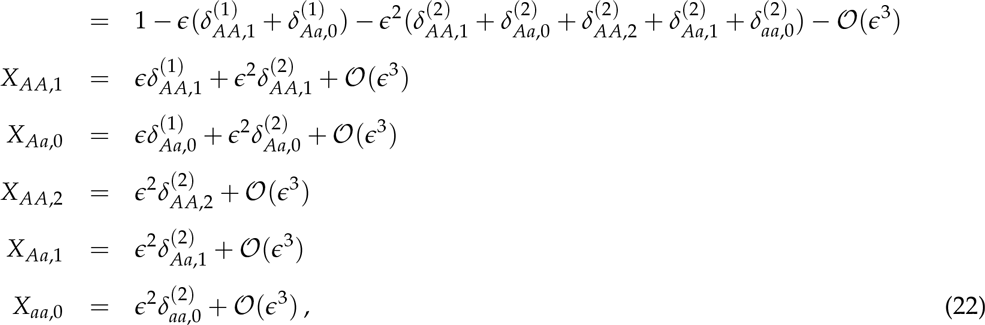
which implies that 
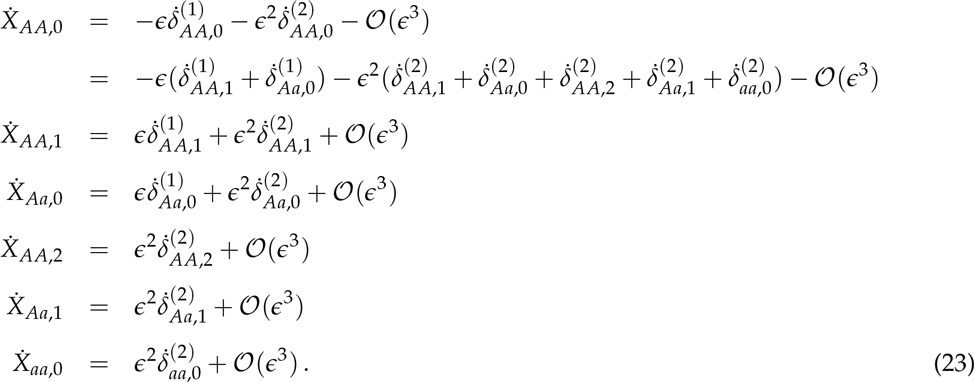

Substituting equation 22 into equation 15, and keeping terms only up to order ɛ^2^, gives 
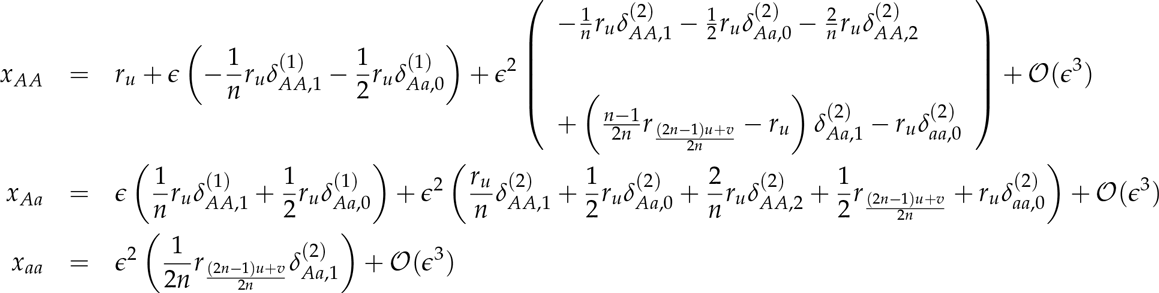

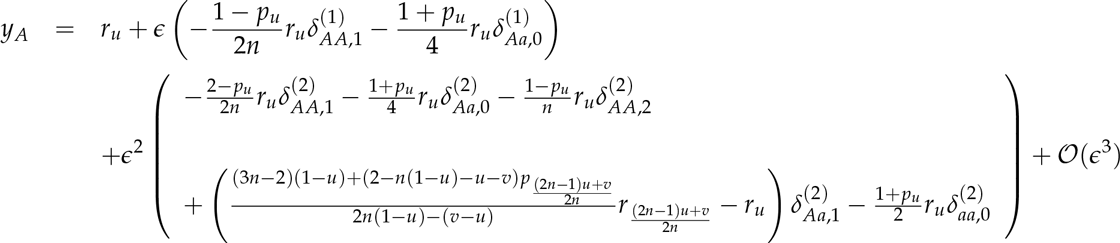

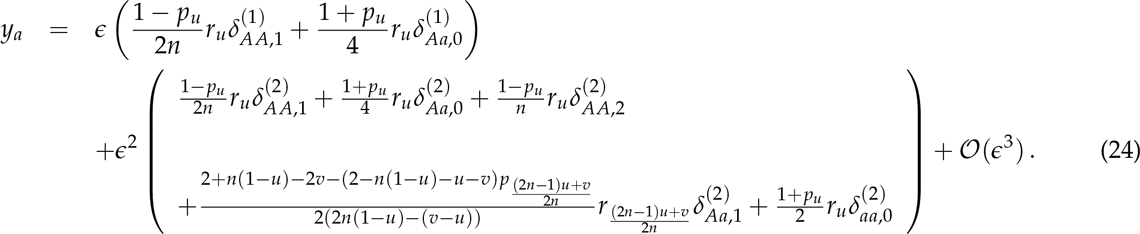

Substituting equations 12, 23, and 24 into equation 21 and discarding powers of ɛ^2^ or higher gives, in matrix form, 
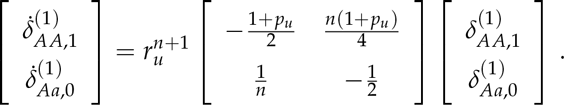

The dominant eigenvalue is 0, and its corresponding eigenvector is 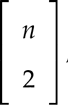, which gives 
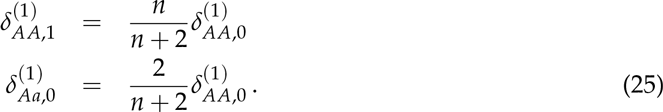

(In other words, this tells us how to “distribute” the first-order perturbation to *X_AA_*,_0_ over the first-order perturbations to *X_AA,1_* and *X_Aa,0_*.)

Substituting equations 12, 23, 24, and 25 into equation 21, and keeping terms up to order ɛ^2^, gives 
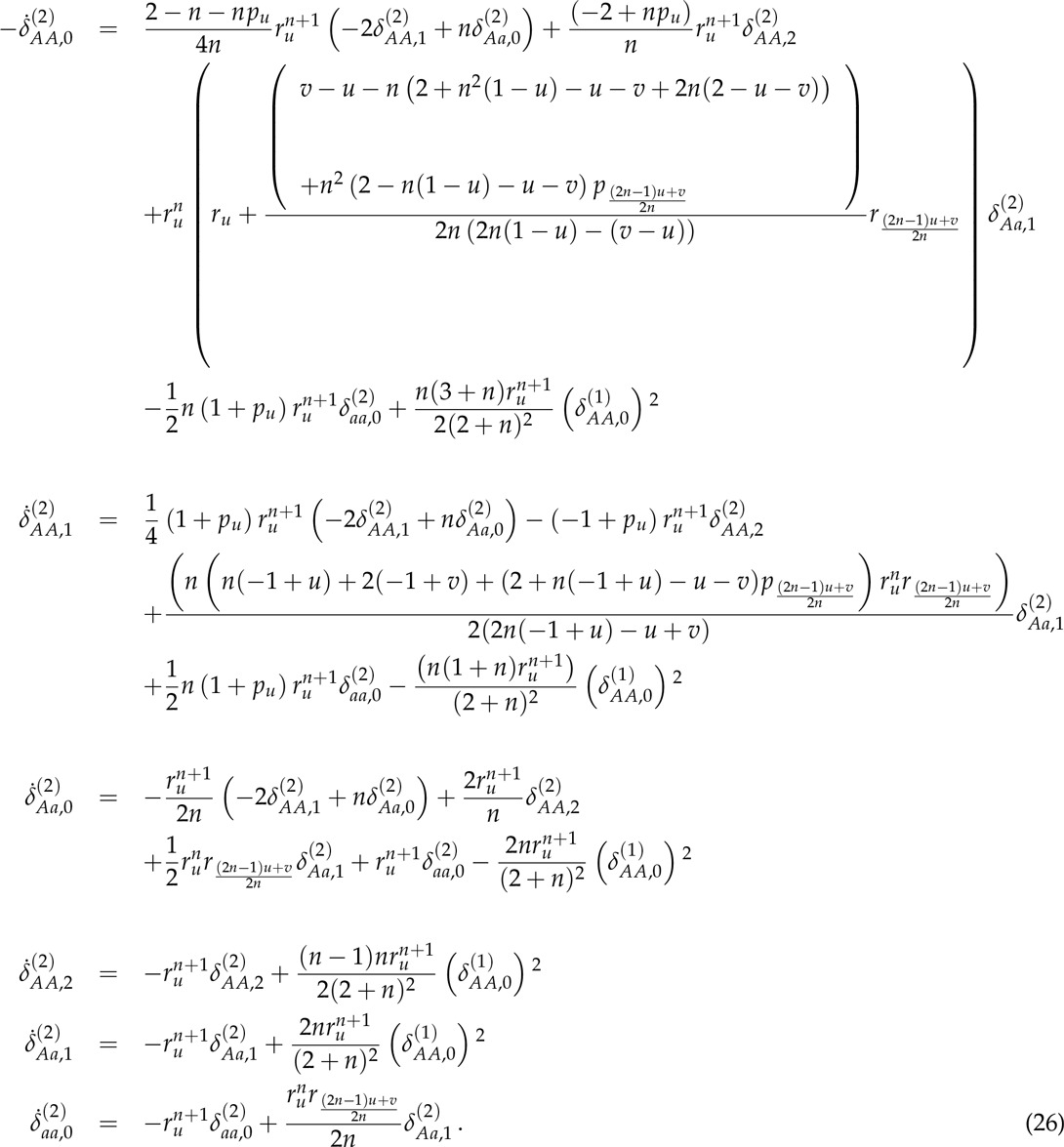

Now, each of these equations must be solved.

The equation for 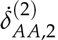 can be directly integrated, yielding: 
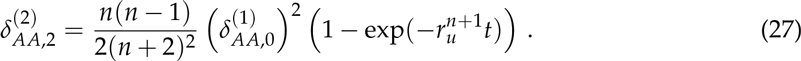

The same can be done for 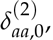, yielding: 
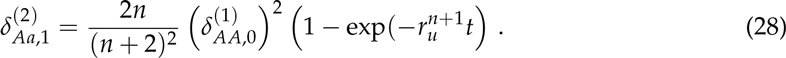

Equation 28 can be used to solve for 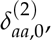, yielding: 
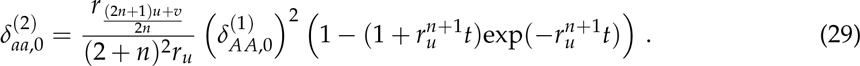

The equations for 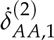 and 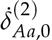 can be manipulated to yield 
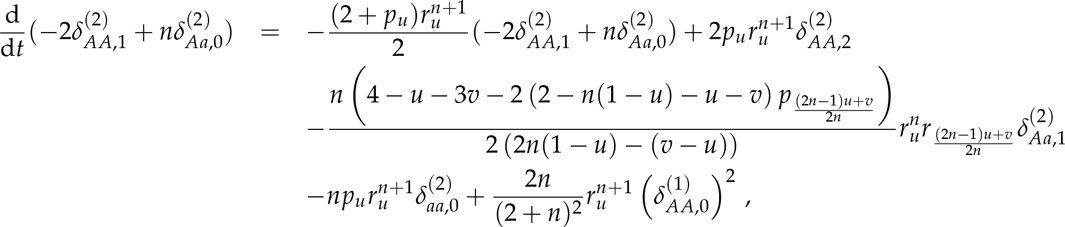
which can be integrated to give 
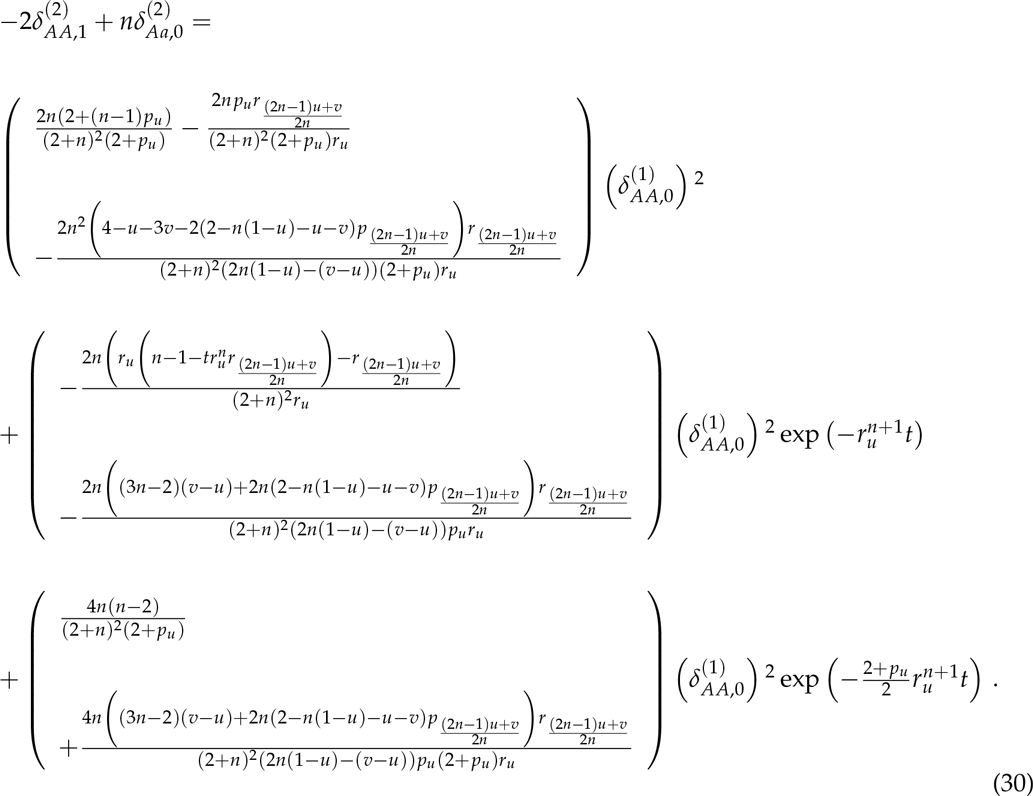

We solve for 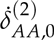 by substituting equations 27-30 into equation 26. In doing so, we permit t to become relatively large, such that all the time-dependent terms in equations 27-30 approach zero. Accordingly, the sign of 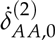 tells us that the mutant sterility allele will invade if: 
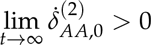

That is, after substitution and simplification, a recessive sterility allele with penetrance *v* will invade a population monomorphic for sterility with penetrance *u* if 
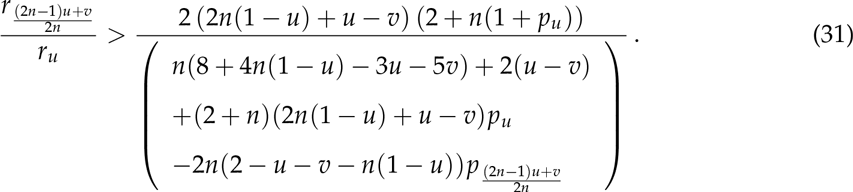

## Appendix B: Kin-selection analysis

Here, we develop a general model of the evolution of wholly or partly non-reproductive workers using standard kin selection methodology (Taylor & Frank 1996, Frank 1998). In this model, a mated queen founds a colony by producing an initial brood of females and/or males. Depending on the model scenario, first-brood females may either mate with first-brood males—from their own or from a different colony—or remain unmated. Then, according to the level of worker sterility z, a focal first-brood female (*i.e.*, a worker) invests a proportion of her resources into helping to raise the colony’s next brood—which consists partly of queen-produced offspring (queen-laid females, notated f, and queen-laid males, notated m) and partly of worker-produced offspring (worker-laid females, notated φ, and worker-laid males, notated μ)—and a proportion of her resources into producing her own offspring. Individuals of the second brood disperse and mate, with each female mating with n males, and mated females then found new patches, restarting the cycle.

In this model, we denote a focal worker’s sterility by Z, the average sterility on a focal patch by z, and the average sterility in the population by 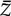. A focal queen’s sex ratio strategy for her second brood is denoted by x, and the average sex ratio strategy among all queens in the population is denoted by 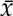. The production of queen-laid second-brood females on a focal patch is *f* = *f* (z, x); the production of queen-laid second-brood males on a focal patch is *m* = *m*(*z*, *x*); the production of worker-laid females by a focal worker is *ϕ* = ϕ(Z, z, *x*); and the production of worker-laid males by a focal worker is *μ* = μ(Z,z, *x*). We denote by 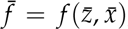, 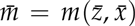, 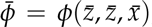, and 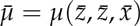 the population-average production of each of these four classes, respectively, and by 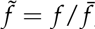, 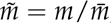, 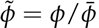, and 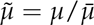 the relative production of each of these four classes.

For a gene increasing worker sterility to spread, its carriers, on average, should leave more descendants than other members of the population. Accordingly, natural selection will favour an increase in worker sterility, z, when 
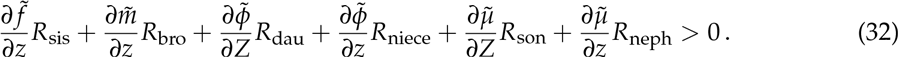

Above, R_sis_, R_bro_, R_dau_, R_niece_, R_son_, and R_neph_ are the (life-for-life) relatedness between a focal female worker and her sister, brother, daughter, niece, son, and nephew, respectively, and all derivatives are evaluated at 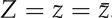.

Each term on the left-hand side of condition 32 captures how a small increase in worker sterility impacts upon the fitness of different individuals in the population, weighted by the life-for-life relatedness between those individuals and a focal worker, which combines both (i) the reproductive value of those individuals (*i.e.*, their capacity for projecting genes into future generations) and (ii) the extent to which those individuals themselves carry the gene increasing worker sterility. Alternatively, each term can be read as an inclusive-fitness effect experienced by a focal worker who gives up reproduction to become sterile. These interpretations are mathematically equivalent, but we focus on the inclusive-fitness interpretation here, as it is conceptually simpler.

Similarly, natural selection will favour an increase in the queen’s sex allocation strategy (her investment in daughters), *x*, when 
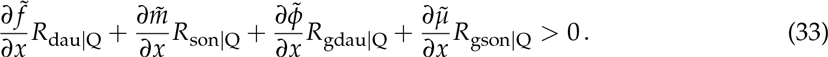

Above, R_dau|Q_ is the relatedness between a focal queen and her daughter, R_son_|Q is the relatedness between a focal queen and her son, R_gdau|Q_ is the relatedness between a focal female and her granddaughter (her daughter’s daughter), R_gson|Q_ is the relatedness between a focal female and her grandson (her daughter’s son), and all derivatives are evaluated at 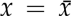. Each term on the left-hand side of condition 33 captures how a small increase in the queen’s investment in daughters, as opposed to sons, impacts upon the fitness of different individuals in the population; alternatively, each term can be read as an inclusive-fitness effect experienced by a focal queen who gives up one of her sons to raise an extra daughter.

For scenario *A*, the production of queen-laid females, queen-laid males, worker-laid females, and worker-laid males is *f* = xr_z_, *m* = (1 — x)*r*_z_*p*_z_, *ϕ* = 0, and 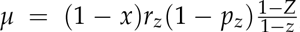, respectively. For scenario B, we use *f* = *x*_*r*z*p*z_, *m* = (1 — x)*r*_z_*p*_z_, ϕ = 0 and 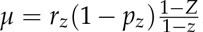. For scenario C, we use *f* = xr_z_p_z_, *m* = (1 — x)r_z_p_z_, 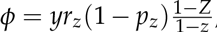,and 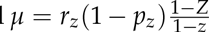. And for scenario D, we use *f* = x(z + sz^2^), *m* = (1 — x)(z + sz^2^), *ϕ* = *y*(1 — Z)(1 — c), and *μ* = (1 − *y*)(1 — Z)(1 — *c*). Substituting these definitions into conditions 32 and 33 recovers conditions 5-10 above.

## Relatedness calculations

The life-for-life relatedness of individual A to individual B is 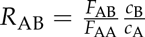, where FAB is the consanguinity of individual A and individual B, F_AA_ is the consanguinity of individual A to herself, *c*_B_ is the class reproductive value of individual B, and *c*_A_ is the class reproductive value of individual A (Bulmer 1994). Note that since individual A is always the same individual within a given condition above, we can instead use R_AB_ = F_ABcB_ or any multiple thereof without affecting the resulting conditions.

Accordingly, consanguinities needed for the conditions above can be found in Table 1. The consanguinities for a female worker under claustral inbreeding are obtained by first calculating the coefficient of inbreeding for a foundress in this mating system (the probability that her two genes at a given locus are identical by descent). Suppose that a juvenile is foundress-laid with probability Q, and soldier-laid with probability 1 — Q. If foundress-laid, her coefficient of consanguinity is zero, because patch founders are unrelated. If worker-laid, then her paternally-inherited gene comes from her grandmother, and her maternally-inherited gene comes, with equal probability, either from her grandfather—who is unrelated to her grandmother—or from her grandmother; in the latter case, her two genes are either copies of the “same” gene in her grandmother, in which case they are identical by descent with probability 1, or are copies of “different” genes from her grandmother, in which case they are identical by descent with probability G, where G is the ju-venile’s grandmother’s coefficient of inbreeding. That is, overall, the probability that these two genes are identical by descent is 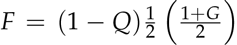, and at equilibrium, G = F, which gives 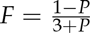. A similar argument gives the same result under diploidy.

**Table 1:** Consanguinities used in inclusive-fitness models.

| For outbreeders |  |  |  |
| --- | --- | --- | --- |
| Relationship | Notation | Haplodiploidy | Diploidy |
| female to daughter | $F_{\text{dau}}$ | $\frac{1}{4}$ | $\frac{1}{4}$ |
| female to son | $F_{\text{son}}$ | $\frac{1}{2}$ | $\frac{1}{4}$ |
| female to sister | $F_{\text{sis}}$ | $\frac{2+n}{8n}$ | $\frac{1+n}{8n}$ |
| female to brother | $F_{\text{bro}}$ | $\frac{1}{4}$ | $\frac{1+n}{8n}$ |
| female to niece | $F_{\text{niece}}$ | $\frac{2+n}{16n}$ | $\frac{1+n}{16n}$ |
| female to nephew | $F_{\text{neph}}$ | $\frac{2+n}{8n}$ | $\frac{1+n}{16n}$ |
| female to daughter's daughter | $F_{\text{gdau}}$ | $\frac{1}{8}$ | $\frac{1}{8}$ |
| female to daughter's son | $F_{\text{gson}}$ | $\frac{1}{4}$ | $\frac{1}{8}$ |
| For claustral inbreeders |  |  |  |
| Relationship | Notation | Haplodiploidy | Diploidy |
| female worker to daughter | $F_{\text{dau\vert c}}$ | $\frac{5+Q}{4(3+Q)}$ | $\frac{11+Q}{8(3+Q)}$ |
| female worker to son | $F_{\text{son\vert c}}$ | $\frac{1}{2}$ | $\frac{11+Q}{8(3+Q)}$ |
| female worker to sister | $F_{\text{sis\vert c}}$ | $\frac{3+2n+Q}{4n(3+Q)}$ | $\frac{1+n}{2n(3+Q)}$ |
| female worker to brother | $F_{\text{bro\vert c}}$ | $\frac{1}{3+Q}$ | $\frac{1+n}{2n(3+Q)}$ |
| female worker to niece | $F_{\text{niece\vert c}}$ | $\frac{3+6n+Q}{8n(3+Q)}$ | $\frac{1+n}{2n(3+Q)}$ |

## Class reproductive values

To determine the class reproductive value of each of the four juvenile classes (queen-laid females, class f; queen-laid males, class φ; worker-laid females, class 9; and worker-laid males, class μ), we first solve for the total reproductive value of all females, C,_F_ = C_f_ + C_φ_, and the total reproductive value of all males, C_M_ = *c*_m_ + *c*_μ_. Defining 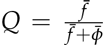 as the probability that a random female is queen-laid, and 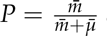 as the probability that a random male is queen-laid, note that a random female inherits half of her genes from a female in the previous census if she is queen-laid, and three quarters of her genes from a female in the previous census if she is worker-laid; and a random male inherits all his genes from a female in the previous census if he is queen-laid, and half of his genes from a female in the previous census if he is worker-laid. Hence, the recurrence relation 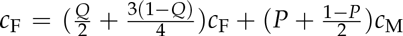, with the constraint that C_M_ = 1 — C_F_, can be solved to give 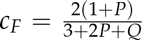 and 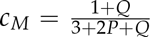. Since an individual’s mating success is not affected by whether they are queen-or worker-laid, we have C_f_ = Q_*c*F_, *c*_φ_ = (1 — Q)*c*_F_, C_m_ = *P*_*c*m_, and C_μ_ = (1 — P)*c*_M_, which, overall, gives

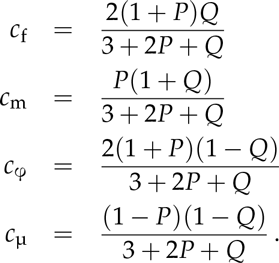

When all second-brood juveniles are queen-laid (*P* = Q = 1), this yields the expected result that C_f_ = 2/3, C_m_ = 1/3, C_φ_ = 0, and C_μ_ = 0; when all second-brood juveniles are worker-laid (P = Q = 0), this yields the expected result that C_f_ = 0, *c*_m_ = 0, *c*_φ_ = 2/3, and *c*_μ_ = 1/3 (Price 1970, Taylor 1996).

It is illustrative to examine a special case. When all second-brood females are queen-laid (Q = 1), this reduces to 
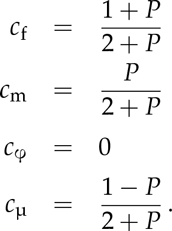

In this case, when *P* = 1, we have the expected result that the total value of juvenile females is 2/3 and the total value of juvenile males is 1/3, because of the usual asymmetries of haplodiploidy. But when *P* = 0, the total value of juvenile females is 1/2 and the total value of juvenile males is 1/2. This is because new juvenile females get half their genes from their mother and half from their father, while new juvenile males are parthenogenetically produced by worker females, and so ul-timately get half their genes from their mother’s mother and half their genes from their mother’s father. In this way, juvenile females and males have an equal share in producing the next genera-tion of juveniles (*cf*. Boomsma & Grafen 1991).

## References

Boomsma, J.J. (2007). Kin selection versus sexual selection: why the ends do not meet. Curr. Biol., 17, R673–R683.

Boomsma, J.J. (2009). Lifetime monogamy and the evolution of eusociality. Phil. T-ans. R. Soc. B, 364, 3191–3207.

Boomsma, J.J. (2013). Beyond promiscuity: mate-choice commitments in social breeding. Phil. Trans. R. Soc. B, 368, 20120050.

Boomsma, J.J. & Grafen, A. (1991). Colony-level sex ratio selection in the eusocial Hymenoptera. J. Evol. Biol., 3, 383–407.

Chapman, T.W., Kranz, B.D., Bejah, K.L., Morris, D.C., Schwarz, M.P. & Crespi, B.J. (2002). The evolution of soldier reproduction in social thrips. Behav. Ecol., 13, 519–525.

Charlesworth, B. (1978). Some models of the evolution of altruistic behaviour between siblings. J. Theor. Biol., 72, 297–319.

Cornwallis, C.K., West, S.A., Davis, K.E. & Griffin, A.S. (2010). Promiscuity and the evolutionary transition to complex societies. Nature, 466,969–972.

Davies, N.G., Ross, L. & Gardner, A. (2016). The ecology of sex explains patterns of helping in arthropod societies. Ecol. Lett., doi: 10.1111/ele.12621

Eshel, I. (1983). Evolutionary and continuous stability. J. Theor. Biol., 103, 99–111.

Frank, S.A. (1998). Foundations of Social Evolution. Princeton University Press, Princeton.

Hamilton, W.D. (1964). The genetical evolution of social behaviour, I & II. J. Theor. Biol., 7,1–52.

Hamilton, W.D. (1972). Altruism and related phenomena, mainly in social insects. Annu. Rev. Ecol. Syst., 3,193–232.

Hughes, W.O.H., Oldroyd, B.P., Beekman, M. & Ratnieks, F.L.W. (2008). Ancestral monogamy shows kin selection is key to the evolution of eusociality. Science, 320, 1213–1216.

Hammerstein, P. (1996). Darwinian adaptation, population genetics and the streetcar theory of evolution. J. Math. Biol., 34, 511–532.

Lukas, D. & Clutton-Brock, T. (2012). Cooperative breeding and monogamy in mammaliansocieties. Proc. R. Soc. Lond. B, 279, 2151–2156.

Maynard Smith, J. & Price, G.R. (1973). The logic of animal conflict. Nature, 246,15–18.

Olejarz, J.W., Allen, B., Veller, C. & Nowak, M.A. (2015). The evolution of non-reproductive workers in insect colonies with haplodiploid genetics. eLIFE, 4, e08918.

Olejarz, J.W., Allen, B., Veller, C., Gadagkar, R & Nowak, M.A. (2016). Evolution of worker policing. J. Theor. Biol., 399,103–116.

Price, G.R. (1970). Selection and covariance. Nature, 227, 520–521.

Ratnieks, F.L.W. (1988). Reproductive harmony via mutual policing by workers in eusocial Hymenoptera. Am. Nat., 132, 217–236.

Ratnieks, F.L.W. & Visscher, P.K. (1989). Worker policing in the honeybee. Nature, 342, 796–797.

Ratnieks, F.L.W., Foster, K.R. & Wenseleers, T. (2006). Conflict resolution in insect societies. Annu. Rev. Entomol., 51,581–608.

Ronai, I., Vergoz, V. & Oldroyd, B.P. (2016). The mechanistic, genetic, and evolutionary basis of worker sterility in the social Hymenoptera. Adv. Stud. Behav., 48, 251–317.

Taylor, P.D. (1996). Inclusive fitness arguments in genetic models of behaviour. J. Math. Biol., 34, 654–674.

Taylor, P.D. & Frank, S.A. (1996). How to make a kin selection model. J. Theor. Biol., 180, 27–37.

Wenseleers, T. & Ratnieks, F.L.W. (2006a). Comparative analysis of worker reproduction and policing in eusocial Hymenoptera supports relatedness theory. Am. Nat., 168, E163–E179.

Wenseleers, T. & Ratnieks, F.L.W. (2006b). Enforced altruism in insect societies. Nature, 444, 50.

